# Species-specific Rates of Fatty Acid Metabolism Set the Scale of Temporal Patterning of Corticogenesis through Protein Acetylation Dynamics

**DOI:** 10.1101/2025.08.27.672586

**Authors:** Ryohei Iwata, Isabel M. Gallego-Lopez, Emir Erkol, Ridha Limame, Anke Vandekeere, Nathan Benac, Benita Turner-Bridger, Mélanie Planque, Martyna Ditkowska, Vera Lein, Franck Maurinot, Nicolas Peredo, Jochen Lamote, Antonela Bonafina, Pieter Vermeersch, Laurent Nguyen, Stein Aerts, Sarah-Maria Fendt, Pierre Vanderhaeghen

**Affiliations:** VIB Center for Brain & Disease Research, 3000 Leuven, Belgium; KU Leuven, Department of Neurosciences & Leuven Brain Institute, 3000 Leuven, Belgium; Laboratory of Cellular Metabolism and Metabolic Regulation, VIB Center for Cancer Biology, VIB, Herestraat 49, 3000 Leuven, Belgium; Laboratory of Cellular Metabolism and Metabolic Regulation, Department of Oncology, KU Leuven and Leuven Cancer Institute (LKI), Herestraat 49, 3000 Leuven, Belgium; Metabolomics Expertise Center, Center for Cancer Biology, VIB, KU Leuven, 3000 Leuven, Belgium; VIB Bioimaging Core Leuven, VIB Technologies, Center for Brain and Disease Research, Leuven, Belgium; VIB Bioimaging Core Leven, Department of Neurosciences, KU Leuven, Leuven Belgium; VIB Flow Core Leuven, VIB Technologies, VIB, Leuven, Belgium; VIB-KU Leuven Center for Cancer Biology, Department of Oncology, Biomedical Sciences Group, KU Leuven, Belgium; Laboratory of Molecular Regulation of Neurogenesis, GIGA Institute, University of Liège, Liège 4000, Belgium; Department of Cardiovascular Sciences, KU Leuven, Leuven, Belgium, and Department of Laboratory Medicine, University Hospitals Leuven, Leuven, Belgium; WELBIO department, WEL Research Institute, avenue Pasteur, 6, 1300 Wavre, Belgium

**Author notes:** These authors contributed equally to this work (Ryohei Iwata, Isabel M. Gallego-Lopez, Emir Erkol).

## Abstract

Developmental processes display temporal differences across species, leading to divergence in organ size and composition. In the cerebral cortex, neurons of diverse identities are generated sequentially through a temporal patterning mechanism conserved throughout mammals. This corticogenesis process is considerably prolonged in the human species, leading to increased brain size and complexity, but the underlying molecular mechanisms remain largely unknown. Here we found that human cortical progenitors displayed lower levels of fatty acid oxidation than their mouse counterparts, in line with their protracted pattern. Treatments that enhance mitochondrial fatty acid oxidation (FAO) accelerated the development of human cortical organoids, including faster progression of neural progenitor cell fate and precocious generation of late-born neurons and glia. FAO accelerated temporal patterning through increased Acetyl-CoA-dependent protein acetylation, including on specific histone transcriptional marks. Thus, species-specific metabolic rates regulate the turnover of post-translation modifications to set the scale of temporal gene regulatory networks of corticogenesis.

## Introduction

Like choreography, development consists of a highly ordered suite of steps and transitions. In the mammalian cerebral cortex, cortical neural progenitor/stem cells called radial glial cells (RGCs) change competence gradually over time, to generate sequentially distinct types of neurons forming the six cortical layers, after which RGCs lose the ability to generate neurons and start producing astroglial cells (*1*, *2*). This process known as temporal patterning of neuronal specification is highly conserved across mammals, including at the level of the underlying gene regulatory mechanisms (*1–3*). However, it is considerably prolonged in the human cortex, lasting 3-4 months instead of 2 months in the macaque, and one week in the mouse species (Fig. 1A). This timing factor is thought to be crucial for increasing cortical size and complexity (*4*, *5*), and it is largely retained by cortical cells cultured from mouse, macaque, and human pluripotent stem cell (PSC), whether in 2D cultures or 3D organoid models (*6–8*), but the underlying mechanisms remain largely unknown.

**Fig. 1.**
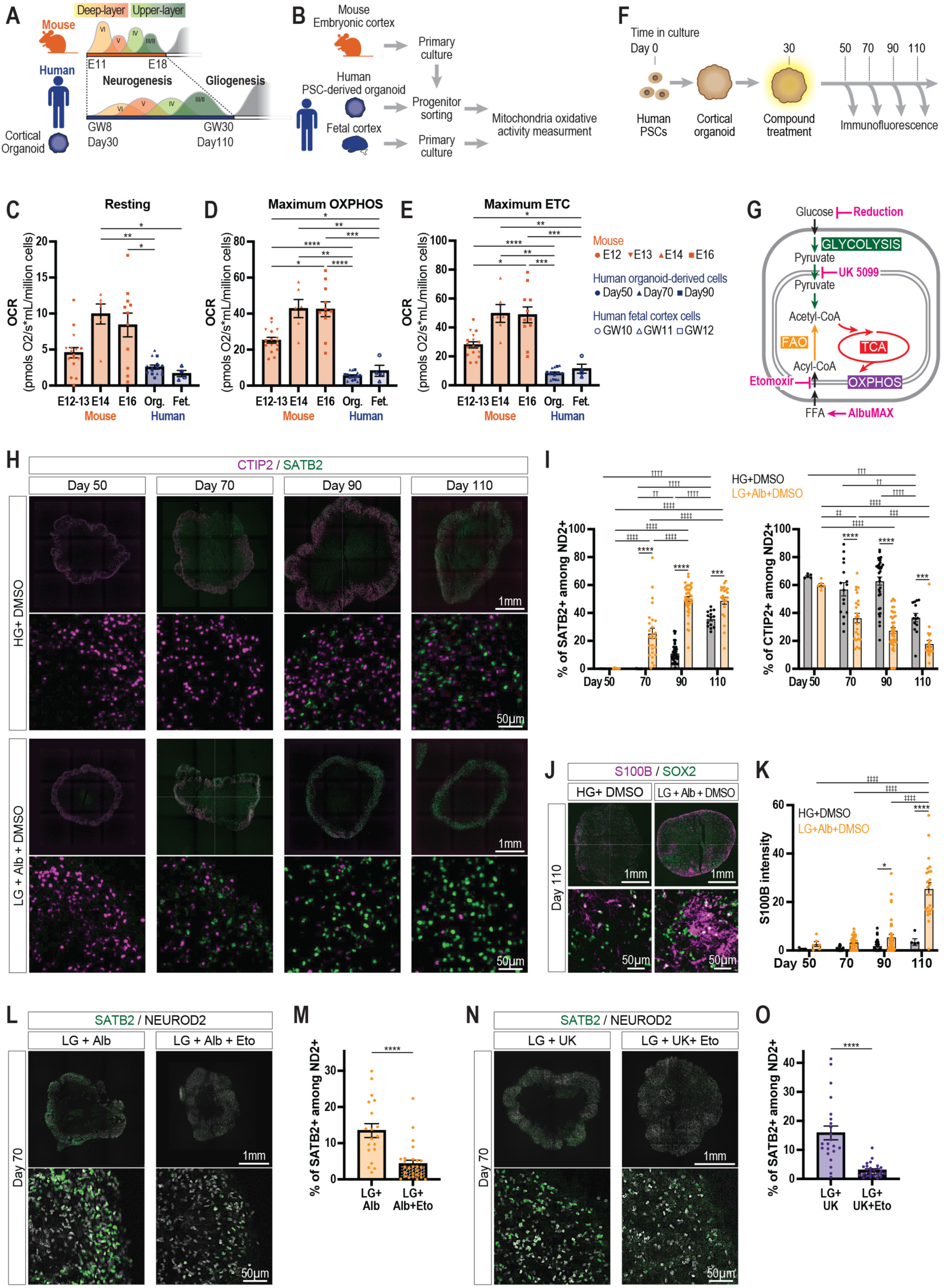
Mitochondria oxidative activity in cortical progenitors displays interspecies differences and influences the timescale of cortical neurogenesis. (A) Schematic of the timeline of temporal patterning in mouse and human developing cortex and cortical organoid models. (B) Schematic of experimental flow to measure mitochondria oxidative activity in mouse and human cortical cells using oxygraph. (C-E) Quantification of oxygen consumption rate (OCR) at different developmental stage cortical progenitors. Each data point represents an individual experiment. Mouse RGCs (derived from embryonic days (E)12, 13, 14, 16. N=9, 7, 6, 11 experiments). Human organoid-derived RGCs (derived from differentiation day 50, 70, 90. N=4, 6, 5 different batches respetively). Human fetal tissue-derived cells (N=4 GW0-12). Dunnett’s or Dunn’s multiple comparisons test. (C) Resting status OCR. (D) Maximum oxidative phosphorylation (OXPHOS) capacity under coupled conditions. (E) Maximum electron transport chain (ETC) capacity under uncoupled condition. (F) Schematic of timeline for human cortical organoid experiments (G) Metabolic pathways targeted by indicated chemical compounds. FAO: Fatty acid oxidation. Tricarboxylic acid cycle: TCA. (H) Representative images of cortical organoid stained for SATB2 and CTIP2. Control HG: High glucose (25 mM) with DMSO (HG+DMSO) (top), Alb: Low glucose (2.5 mM) with 0.5% AlbuMAX and DMSO (LG+Alb+DMSO) (bottom). (I) Quantification of the proportion of (Left) SATB2-positive cells and (Right) CTIP2-positive cells among NEUROD2-positive cells. Each data point represents an individual organoid. HG+DMSO: differentiation day 50, 70, 90, 110. N=5, 17, 36, 14 organoids from six independent differentiation batches. LG+Alb+DMSO: differentiation day 50, 70, 90, 110. N=6, 26, 40, 23 organoids from six independent differentiation batches. Tukey’s multiple comparisons test. (J) Representative images of cortical organoids. S100B is used for labelling astroglial cells. (K) Quantification of mean signal intensity of S100B from each organoid. Each data point represents an individual organoid. HG+DMSO: differentiation day 50, 70, 90, 110. N=4, 24, 32, 5 organoids from ten independent differentiation batches. LG+Alb+DMSO: differentiation day 50, 70, 90, 110. N=6, 31, 41, 23 organoids from ten independent differentiation batches. Tukey’s multiple comparisons test. (L) Representative images of cortical organoid at day 70 for SATB2. Low glucose with 0.5% AlbuMAX and DMSO. Low glucose with 0.5% AlbuMAX and 50µM Etomoxir. (M) Quantification of the proportion of SATB2-positive cells. Each data point represents an individual organoid. Low glucose with 0.5% AlbuMAX and DMSO (LG+Alb): N=19 organoids from three independent differentiation batches. Low glucose with 0.5% AlbuMAX and 50µM Etomoxir (LG+Alb+Eto): N=27 organoids from three independent differentiation batches. Mann-Whitney test. (N) Representative images of cortical organoid at day 70 for SATB2. Low glucose with 1.25µM UK 5099 (LG+UK). Low glucose with 1.25µM UK 5099 and 50µM Etomoxir (LG+UK+Eto). (O) Quantification of the proportion of SATB2-positive cells. Each data point represents an individual organoid. Low glucose with 1.25µM UK 5099 (LG+UK): N=22 organoids from three independent differentiation batches. Low glucose with 1.25µM UK 5099 and 50µM Etomoxir (LG+UK+Eto): N=25 organoids from three independent differentiation batches. Mann-Whitney test. * † ‡P < 0.05, ** †† ‡‡P < 0.01, *** ††† ‡‡‡P < 0.001, **** †††† ‡‡‡‡P < 0.0001.

Species-specific differences in developmental tempo have been studied recently in several cellular contexts, from somitic oscillations to motor neuron generation and cortical neuron maturation (*9*– *15*). In each of these systems, the underlying mechanisms have been linked to species differences in various global cellular processes, including rates of RNA and protein turnover, intermediate metabolism, or chromatin remodeling. These data point to global regulators of developmental timing acting upstream of the gene regulatory networks specific to each cellular system. However, it remains unclear whether any of the proposed mechanisms apply to every system considered, and if they act in parallel or linked to one another. Moreover, none of these global mechanisms have been tested in the most complex sequence of cell fate transitions characterizing cortical temporal patterning.

Several metabolic pathways, particularly at the mitochondria level, were previously implicated in the regulation of balance and differentiation of neural stem cells, from fly to human species (*16– 29*). But whether and how they contribute to temporal patterning of neuronal specification remains unknown. Here we tested whether and how mitochondria metabolism could be linked to the species-specific extended timescale of cortical temporal patterning in the human species.

### Mitochondria oxidative metabolism is lower in human than in mouse cortical progenitors

As temporal patterning is thought to be largely encoded within cortical progenitors (*1*, *2*, *30*), we first determined whether there were any detectable differences in metabolic activity between human and mouse RGCs. To this end, human and mouse cortical RGCs were purified using magnetic-activated cell sorting (MACS) from in vitro cultures obtained from mouse embryonic cortex (E12-E16) and human PSC-derived cortical organoids (day50-day90), corresponding to cortical neurogenesis (*31*, *32*). This led to cell preparations containing > 70% Pax6-positive RGCs (fig. S1, A to C), which were then analyzed acutely for mitochondrial metabolic activity, using oxygen consumption as a generic read-out. Oxygraphy-based measurements performed on the purified preparations revealed that levels of mitochondria oxygen consumption tended to increase over embryonic time within both species, as previously observed in the mouse cortex (*19*). However, the levels of basal and maximal oxidative phosphorylation and electron transport chain were all consistently higher in mouse than human cortical RGCs, at all stages examined (Fig. 1, C to E and fig. S1, D to I). Importantly, similar species-specific values of oxygen consumption were measured in ex vivo cultures of human fetal brains (GW10-12) (Fig. 1, C to E). These data indicate that human cortical progenitors display lower mitochondrial oxidative rates than their mouse counterparts, in correlation with their prolonged timescale of temporal patterning.

### Increasing mitochondria metabolic activity in human cortical organoids leads to accelerated generation of late-born neurons and glia

We next tested whether the observed species differences in mitochondria metabolic rates could be causally linked to the timescale of temporal patterning of corticogenesis. We first applied to human cortical organoids a metabolic treatment (referred to as LG+Alb+GSK) (fig. S2, A and B) previously found to enhance mitochondria metabolism and maturation speed in human postmitotic neurons (*13*). LG+Alb+GSK consists of low glucose concentration (2.5 mM) (LG, in contrast to high glucose (HG) concentration (25 mM) classically used in neural cell/organoid cultures), a chemical inhibitor of lactate dehydrogenase A (LDHA) (GSK-2837808A (referred to as GSK)), and a mixture of long-chain free fatty acids (FFA) (AlbuMAX referred to as (Alb)) (fig. S2, A and B). As expected, LG+Alb+GSK led to increased mitochondria respiration in human cortical RGCs (fig. S1, J to M). Human cortical organoids were cultured in control conditions (HG+DMSO) or treated with LG+Alb+GSK from early stages of neurogenesis (day 30; counted from day 0 as the start of neural induction from PSC), and analyzed over the entire course of cortical neurogenesis, until the onset of gliogenesis (day 50 to day 110) (fig. S2A). We analyzed fate markers that reflect temporal patterning: CTIP2 (BCL11B), a marker appearing at earlier stages expressed in deep-layer / non-intratelencephalic (DL/non-IT) neurons, , and SATB2, a marker appearing at later stages in intra-telencephalic and upper layer (UL/IT) neurons,. In control conditions, CTIP2- positive neurons constituted the majority of the neuronal population at early stages (day 50), then gradually decreased, while SATB2-positive neurons appeared at later stages (d90-110) (fig. S2, C and D). In contrast, the LG+GSK+Alb treated organoids displayed accelerated increase in the proportion of SATB2-positive neurons (at day 70-90), together with precocious decreased proportion of CTIP2-positive neurons (fig. S2, C and D). S100B, a marker of astrogliogenesis that starts at the end of neurogenesis, was also increased precociously in the LG+GSK+Alb condition (fig. S2, E and F). The same results were obtained in two independent ESC and iPSC lines (fig. S2, G and H). Overall, these data indicate that increased mitochondria metabolic activity leads to faster progression of corticogenesis, leading to precocious onset of expression of late-born-like neuronal and glial fates, normally appearing several weeks later in control conditions.

### FAO is the main metabolic pathway that accelerates human corticogenesis

To determine which specific metabolic pathway was involved in the observed effects, we exposed cortical organoids to single treatments, focusing on the same markers at mid-corticogenesis (day 70) (fig. S3, A to D). LG led to a moderate increase of late fate marker expression, LDH inhibition (GSK) alone had no detectable effect, but Alb condition together with LG led to a maximal effect, similar to the one observed with LG+GSK+Alb (fig. S3B). These data point to FFA as the main driver of the metabolic effects on the tempo of corticogenesis. We next examined in depth the impact of Alb throughout corticogenesis. This revealed a precocious appearance of SATB2+ UL/IT neurons and a precocious decrease in CTIP2+ DL-non-IT neurons (Fig. 1, H and I, and fig. S4). Alb treatment also resulted in precocious onset of gliogenesis, detected with S100B, although less efficiently than together with GSK, suggesting additional effects of LDH inhibition on gliogenesis (Fig. 1, J and K).

Importantly, the effects of FFAs were blocked by a CPT1A inhibitor, Etomoxir, that inhibits the transport of long-chain FFA in mitochondria and thereby prevents fatty oxidation (FAO) (Fig. 1, L and M, and fig. S5, A and B). These data further indicate that the effects of FFA are mostly linked to mitochondria FAO. In line with this, SATB2 expression could also be induced precociously by treatments of the organoids with palmitate, a long-chain FFA that is a common physiological substrate of FAO (fig. S5, E to H).

Interestingly, each of the investigated metabolic treatments resulted in similarly increased levels of oxidative phosphorylation, as measured by oxygraphy (fig. S3, E to H), suggesting that oxygen consumption and oxidative phosphorylation (OXPHOS) *per se* are not the main metabolic outcome required for the effects of FFA on fate changes. In line with this hypothesis, we tested the impact of blockade of pyruvate entry into mitochondria using mitochondria pyruvate carrier (MPC) inhibitor, UK-5099 (referred to as UK) (*33*), under LG condition (Fig. 1G). Remarkably, this led to a strong increase in late neuronal fate markers and increased oxygen consumption (fig. S3, A,B, E to H). We reasoned that this could be explained as UK treatment in presence of low glucose would be expected to lead to increased mitochondria FAO, as previously reported in other systems (*34*). Confirming this hypothesis, UK effect on increased SATB2 was abolished by FAO inhibition (Fig. 1, N and O, and fig. S5, C and D).

These data indicate that FFA-driven FAO rates can act as powerful accelerators of temporal patterning in human cortical organoids.

### FAO accelerates temporal patterning of neuronal specification at the level of RGCs

A hallmark of temporal patterning of neuronal specification is that neurons of distinct identities are born at different time points. While our data so far were consistent with effects on temporal patterning, they could also be interpreted as shifts in expression of the fate markers in the neurons, without actual changes in their timing of generation. To distinguish between these possibilities, we next examined whether the actual birthdate of neurons of specific identities was also temporally shifted in accordance with accelerated temporal patterning. While pulse-chase nuclear labeling is classically used in vivo to determine neuronal birthdate, it remains very difficult to perform for in vitro human corticogenesis, because of long-term neuronal toxicity. We therefore used a method that we previously developed, in which cohorts of neurons generated at the same time using γ-secretase inhibitor DAPT (a Notch inhibitor that increases rates of neurogenesis) are xenotransplantated in the mouse neonatal cortex (*13*, *35*) (Fig. 2). Crucially, in this system the human neurons that are integrated in the mouse cortex are born within a few days prior to the transplantation (*35*), thus allowing to analyze cohorts of neurons born at the same time, and to quantify the outcome of cell fate specification in vivo (Fig. 2A). Human cortical organoids were first cultured in control conditions or treated with FFA from day 30 as above. Then, at various time-points (day 56, 70 and 84), the organoids were dissociated, followed by acute DAPT treatment to synchronize newborn neuron induction coupled with xenotransplantation in the neonatal mouse (postnatal day 0 or 1) cortex, and in vivo examination of neuron fate markers (CTIP2 and SATB2) 14 days later (Fig. 2A). In control conditions, we observed a gradual increase in the proportion of SATB2 expression among the neurons prepared from organoids cultured for longer periods, while the proportion of CTIP2 neurons was decreased accordingly (Fig. 2, B to E). This confirmed that the xenotransplantation method allows the analysis of the timing of fate specification within a cohort of neurons sharing a similar birthdate. Importantly, this temporal patterning was accelerated in the cortical organoids following treatment with Alb, reflected by a precocious increase in the generation of SATB2 neurons and a precocious decrease in the generation of CTIP2 neurons (Fig. 2, B to E). These data suggest that enhanced FAO in cortical organoids leads to an accelerated change in the competence of RGCs to generate distinct type of neurons at different timepoints, thus reflecting a genuine change in temporal patterning, and not merely a switch in expression of fate markers.

**Fig. 2.**
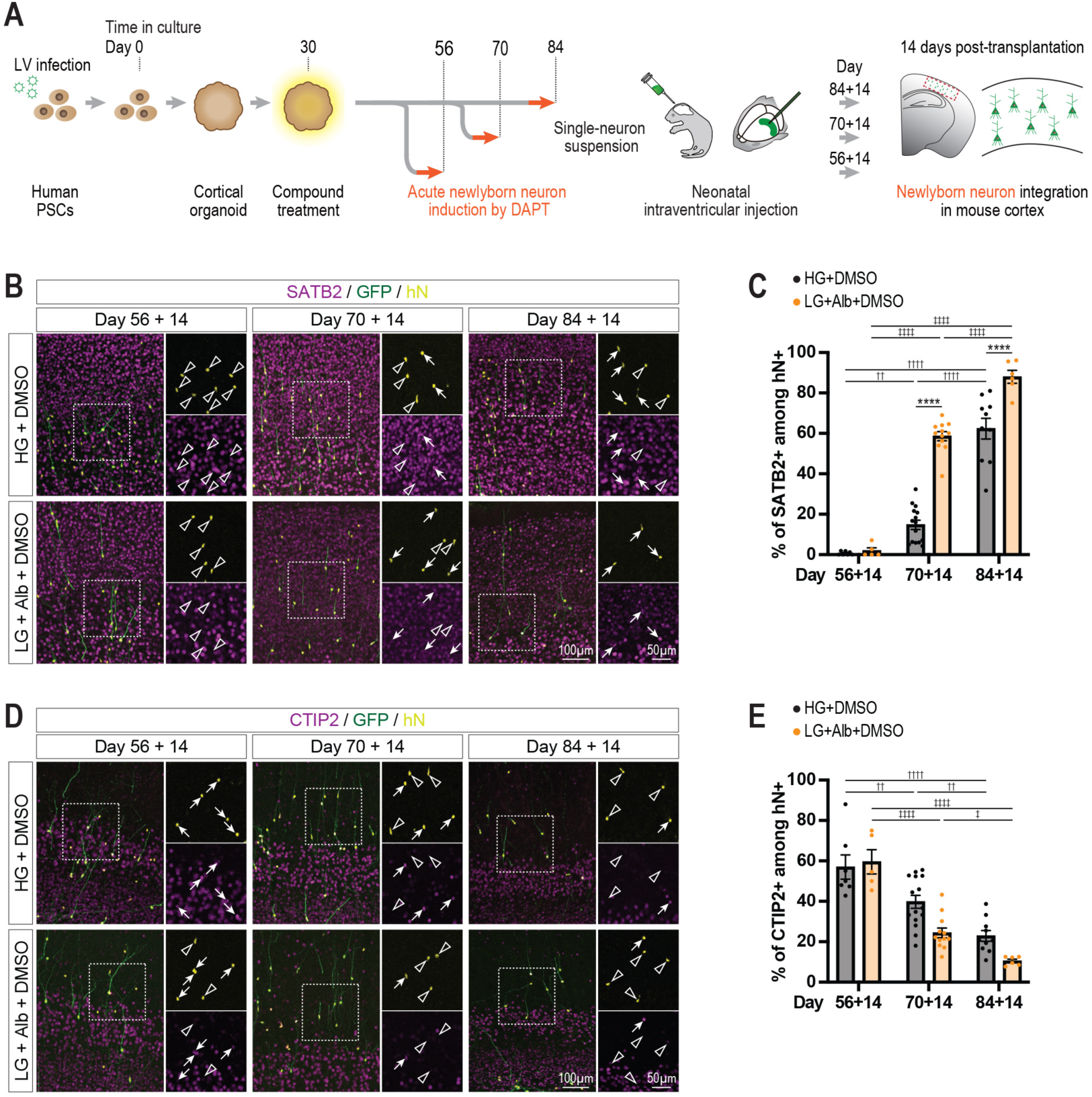
Enhanced mitochondrial fatty acid oxidation leads to precocious generation of late-born neurons. (A) Experimental scheme of xenotransplantation of cortical human organoid-derived newborn neurons into neonatal mouse. (B) Representative images of human cortical neurons at 14 days post-transplantation in the mouse cerebral cortex. Arrows indicate SATB2-positive and arrowheads indicate SATB2-negative cells. (C) Quantification of the proportion of SATB2-positive cells. Each data point represents an individual xenotransplanted mouse. High glucose with DMSO. HG: differentiation day 56, 70, 84, N=7, 15, 10 mice using three independent organoid differentiation batches. Low glucose with 0.5% AlbuMAX and DMSO. LG+Alb: differentiation day 56, 70, 84, N=5, 12, 6 mice using three independent organoid differentiation batches. Tukey’s multiple comparisons test. (D) Representative images of human cortical neurons at 14 days post-transplantation in the mouse cerebral cortex. Arrows indicate CTIP2-positive and arrowheads indicate CTIP2-negative neurons. (E) Quantification of the proportion of CTIP2-positive cells. Each data point represents an individual xenotransplanted mouse. High glucose with DMSO (HG): differentiation day 56, 70, 84, N=7, 15, 10 mice using three independent organoid differentiation batches. Low glucose with 0.5% AlbuMAX and DMSO (LG+Alb): differentiation day 56, 70, 84, N=5, 12, 6 mice using three independent organoid differentiation batches. Tukey’s multiple comparisons test. * † ‡P < 0.05, ** †† ‡‡P < 0.01, *** ††† ‡‡‡P < 0.001, **** †††† ‡‡‡‡P < 0.0001.

### Enhanced FAO accelerates the molecular temporal patterning of cortical RGCs

We next analyzed the impact of FFA on cortical organoids with single-nucleus RNA sequencing (snRNAseq) to determine the cellular and molecular mechanisms underlying its temporal effects (Fig. 3A). Cortical organoids were treated with FFA from the onset of neurogenesis (day 30), and their transcriptome examined at 3 time-points throughout corticogenesis (day 50, 70 and 90). Cell annotations were assigned based on marker genes (fig. S6E) derived from in vivo human and organoid datasets (*36*, *37*) and further confirmed by integration to in vivo fetal data (*38*). This revealed that most of the organoid cells correspond to cortical cells found in vivo at all stages examined in both control and FFA conditions (Fig. 3, B and D and fig. S6, A to D).

**Fig. 3.**
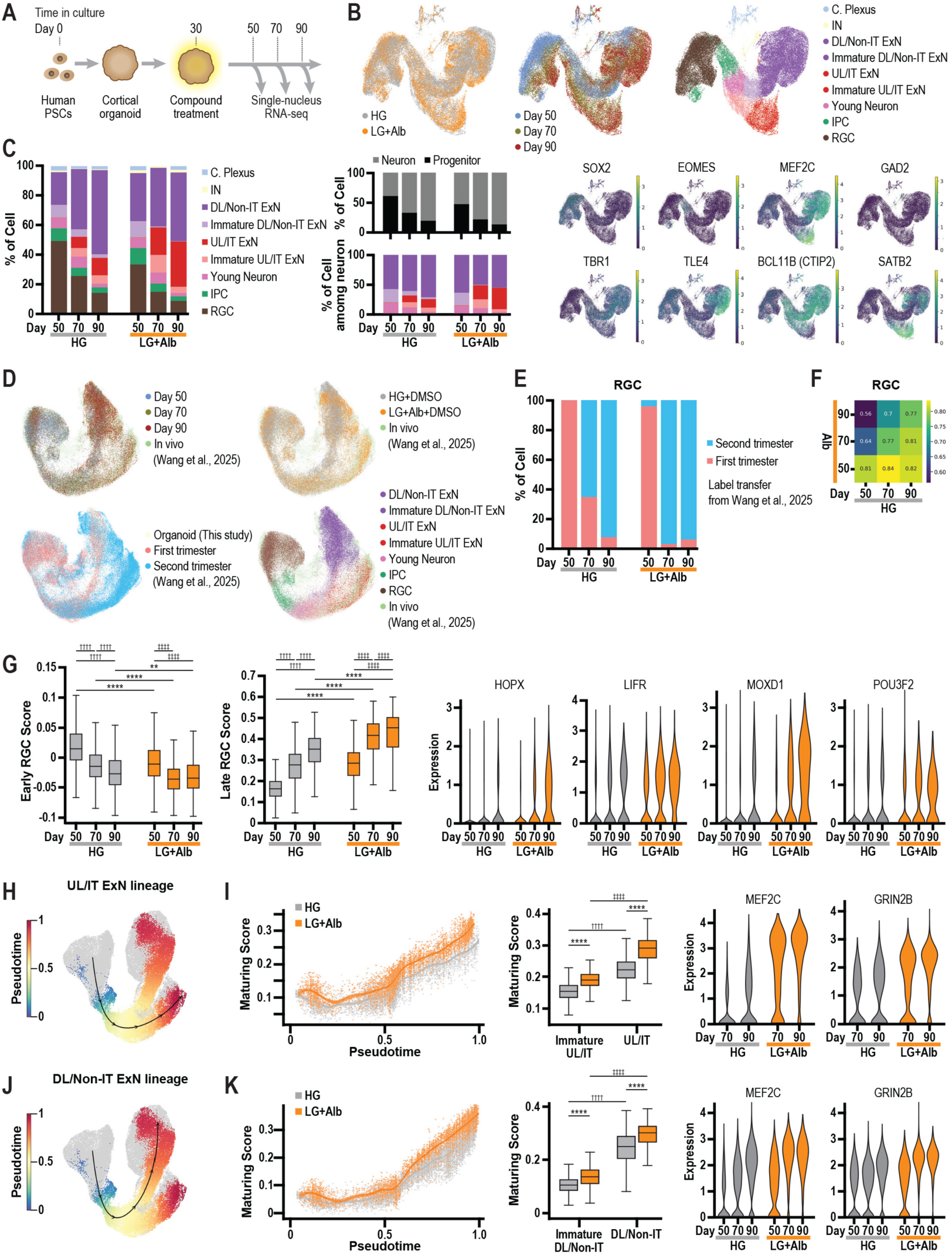
Fatty acid oxidation accelerates temporal patterning in cortical progenitors and maturation in neurons. (A) Schematic of timeline for human cortical organoid cultures and treatments for snRNAseq experiments. (B) UMAP plot showing untreated and treated organoids (Top left), Age of organoids (Top middle), major cell type annotations (Top right) and markers of major cell types (Bottom). (C) Cell type proportions across age and treatments in cortical organoids. (Left) Proportion of neurons and progenitors among all cells. (Right top) Proportion of each neuron subtype among all neurons (Right bottom). (D) UMAP plot of in vivo and organoids showing age in organoids (Top left), treatment in organoids (Top right), in vivo human fetal cortex (Bottom left) and cell types (Bottom right) after Harmony integration. (E) Temporal stage predictions of cortical RGCs using label transfer from in vivo human fetal cortex labels of organoid RGCs. (F) Heatmap of Spearman correlation of RGCs of treated and untreated conditions (G) (Left) Gene set score of RGCs for late RGC gene markers. Mann-Whitney test two-sided with Bonferroni correction. (Right) Violin plot showing normalized expression of differentially expressed canonical late RGC markers. (H) Pseudotime scores of IT excitatory neuron trajectory displayed over UMAP plot. (I) (Left) Generalized additive model (GAM) fit of maturing IT neuron score along control and FFA-treated IT neuron trajectories. (Middle) Boxplot of mature IT score over immature IT and IT neurons across control and Alb treatment. (Right) Normalized expression of differentially expressed fetal cortex neuron maturation markers. (J) Pseudotime scores of DL/Non-IT excitatory neuron trajectory displayed over UMAP plot. (K) (Left) GAM fit of maturing DL/Non-IT neuron scores along control and Alb-treated DL/Non-IT neuron trajectories. (Middle) Boxplot of maturing DL/Non-IT score over immature DL/Non-IT and DL/Non-IT neurons across control and Alb treatment. (Right) Normalized expression of differentially expressed fetal cortex neuron maturation markers. * † ‡P < 0.05, ** †† ‡‡P < 0.01, *** ††† ‡‡‡P < 0.001, **** †††† ‡‡‡‡P < 0.0001.

We first examined cell distribution and identity in control cortical organoids over the 3 experimental time-points. We found a gradual decrease over time in the proportion of RGCs, a parallel increase in neuronal proportion, and most strikingly an increase in the proportion of late born-like UL/IT neurons and conversely a gradual decrease in the proportion of early born-like DL/non-IT neurons, thus reflecting cortical temporal patterning (Fig. 3C). Remarkably, all these temporal changes appeared 3-6 weeks earlier in organoids treated with Alb, extending our observations at the protein level (Fig. 1 and 2), and consistent with a globally precocious temporal patterning of neuronal specification (Fig. 3C).

We next analyzed the transcriptome of RGCs, focusing on changes that underlie temporal patterning. We performed temporal prediction of organoid RGCs using a semi-supervised deep generative model (scANVI) (*39*) implemented within the scArches framework (*40*) trained on in vivo human fetal RGC data from first and second trimester stages (*38*). This revealed that D70 Alb-treated organoids showed an increased proportion of RGCs predicted to be in second trimester stage (Fig. 3E). Moreover, temporal comparison of the transcriptome of RGCs between conditions revealed that the treated RGCs were more similar to untreated RGCs analyzed at later time-points, consistent with acceleration of temporal patterning of Alb-treated RGCs (Fig. 3F). To test this further, we scored RGCs with a gene set corresponding to "early" and "late" RGCs, previously generated using first trimester human fetal brain developmental atlas data (*41*). This analysis revealed an increase in expression of late RGC markers over time, and conversely a decrease in early RGC markers, and this was accelerated in the RGCs from Alb-treated organoids (Fig. 3G and fig. S6F). Accordingly, Alb-treated RGCs displayed faster increase of canonical markers of the RGCs that normally emerge later in human corticogenesis (Fig. 3G, HOPX, LIFR, MOXD1 and POU3F2). Interestingly, these late RGC markers are also also highly enriched outer radial glial cells, a key population of neural stem/progenitor cells uniquely expanded in the human cortex (*42*, *43*). These data indicate that FFA accelerates the temporal progression of RGCs at the level of their whole transcriptome, resulting in precocious specification of late born-like UL/IT cortical neurons.

Finally we examined cortical neuron maturation, which was previously shown to be influenced by mitochondria metabolism (*13*). We first performed temporal analyses on the pseudotime trajectories reconstructed for both UL/IT and DL/Non-IT excitatory neuron lineages (Fig. 3, H and J). We scored cortical neurons in each of the lineage according to a gene set reflecting neuronal maturity. To generate this gene set, we performed a differential expression test between immature DL/non-IT and more mature DL/non-IT neurons in control organoids (Fig. 3B). Genes identified as upregulated constituted the "maturing" gene set and genes identified as downregulated constituted the "immature" gene set. The same analysis was performed for IT neurons. Remarkably, both "maturing" and "immature" gene set score were accelerated in both neuron trajectories in Alb-treated organoids (Fig. 3, I and K, and fig. S6, G and H). Accordingly, Alb-treated neurons displayed faster increase of canonical markers of neuronal maturation (such as MEF2C and GRIN2B) (Fig. 3, I and K) and upregulated expression highly enriched for genes associated with axon, dendrite, and synapse development (Tables S1 and S2).

Overall, these data indicate that increased FFA availability is sufficient to accelerate global timing of gene expression in human cortical organoids, from temporal patterning of neural progenitors to neuronal subtype specification to maturation.

### FAO rates differ between mouse and human RGCs and impact on temporal patterning through Acetyl-CoA-dependent protein acetylation

Given the effects of FAO on the timescale of human corticogenesis, we next tested their relevance on species differences in temporal patterning. We compared FAO rates in mouse and human cortical progenitors purified by MACS as described above (Fig. 4A). Oxygraphy combined with FAO substrates, revealed higher FAO-dependent activity in mouse compared with human RGCs (Fig. 4B, and fig. S7, A and B). Moreover, measurement of lipid uptake using BODIPY-C12 labeling (*44*) on live cortical cells revealed higher uptake rates in mouse than human cortical RGCs (Fig. 4, C to E, and fig. S7, C and D). These data indicate that mouse and human cortical RGCs display species-specific differences in lipid uptake and mitochondria FAO that correlate with the temporal scale of corticogenesis in each species.

**Fig. 4.**
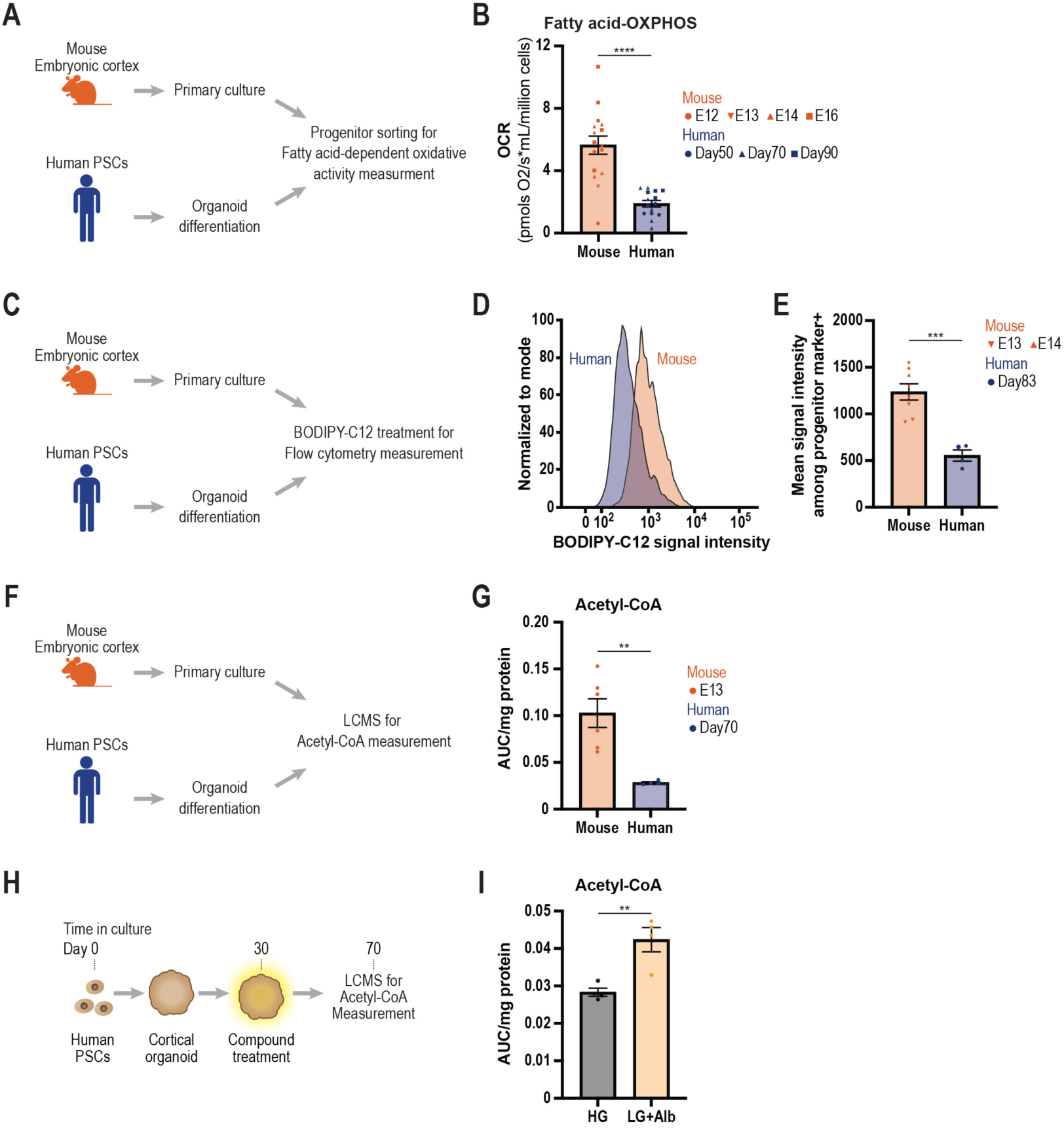
Interspecies differences in FAO rates and Acetyl-CoA. (A) Schematic of experimental flow of fatty acid-dependent mitochondria oxidative (fatty acid-OXPHOS) activity measurement using oxygraphy in cortical RGC progenitors. (B) Quantification of fatty acid treatment-induced OCR. Each data point represents an individual experiment. Mouse embryonic cortex RGC progenitors (derived from embryonic days (E)12, 13, 14, 16. N=4, 1, 6, 5 experiments). Human cortical organoid-derived RGC progenitors (derived from differentiation day 50, 70, 90. N=4, 6, 5 different batches). Unpaired t test. (C) Schematic of experimental flow of BODIPY-C12 uptake activity measurement using flow cytometry. (D) Representative plot of BODIPY intensity in mouse and human cortical progenitors. (E) Quantification of mean signal intensity of BODIPY-C12 of cortical progenitors. Each data point represents an individual experiment. Mouse primary progenitors (derived from embryonic days (E)13, 14. N=6, 2 experiments). Human organoid-derived progenitors (derived from differentiation day 83. N=4 different batches). (F) Schematic of experimental flow of Acetyl-CoA measurement using Liquid chromatography-mass spectrometry (LCMS). (G) Quantification of Acetyl-CoA in mouse and human cortical cells. Each data point represents an individual experiment. Mouse cortical cells (derived from E13. N=6 experiments). Human organoid-derived cortical cells (derived from differentiation day 70. N=4 experiments from two independent differentiation batches). Unpaired t test. AUC: area under the curve. (H) Schematic of experimental flow of Acetyl-CoA measurement in chemical compound-treated cortical organoids. (I) Quantification of Acetyl-CoA in human cortical organoids cultured in HG or LG+Alb conditions. Each data point represents an individual experiment. HG+DMSO: Derived from differentiation day 70. N=4 experiments from two independent differentiation batches. LG+Alb+DMSO: Derived from differentiation day 70. N=4 experiments from two independent differentiation batches. Unpaired t test. **P < 0.01, ***P < 0.001, ****P < 0.0001.

How could FAO metabolism impact global temporal patterns of cell fate during corticogenesis? In many cell types, FAO has been shown to promote cell fate transitions by increasing the levels of Acetyl-CoA, leading to enhanced protein acetylation, including on specific histone marks linked to chromatin remodeling (*45–50*). In these cases, Acetyl-CoA originate mainly from the conversion of mitochondria TCA-derived citrate that is transported outside of mitochondria, followed by its conversion into Acetyl-CoA through ATP citrate lyase (ACLY) (*45*, *51*) (Fig. 5A).

**Fig. 5.**
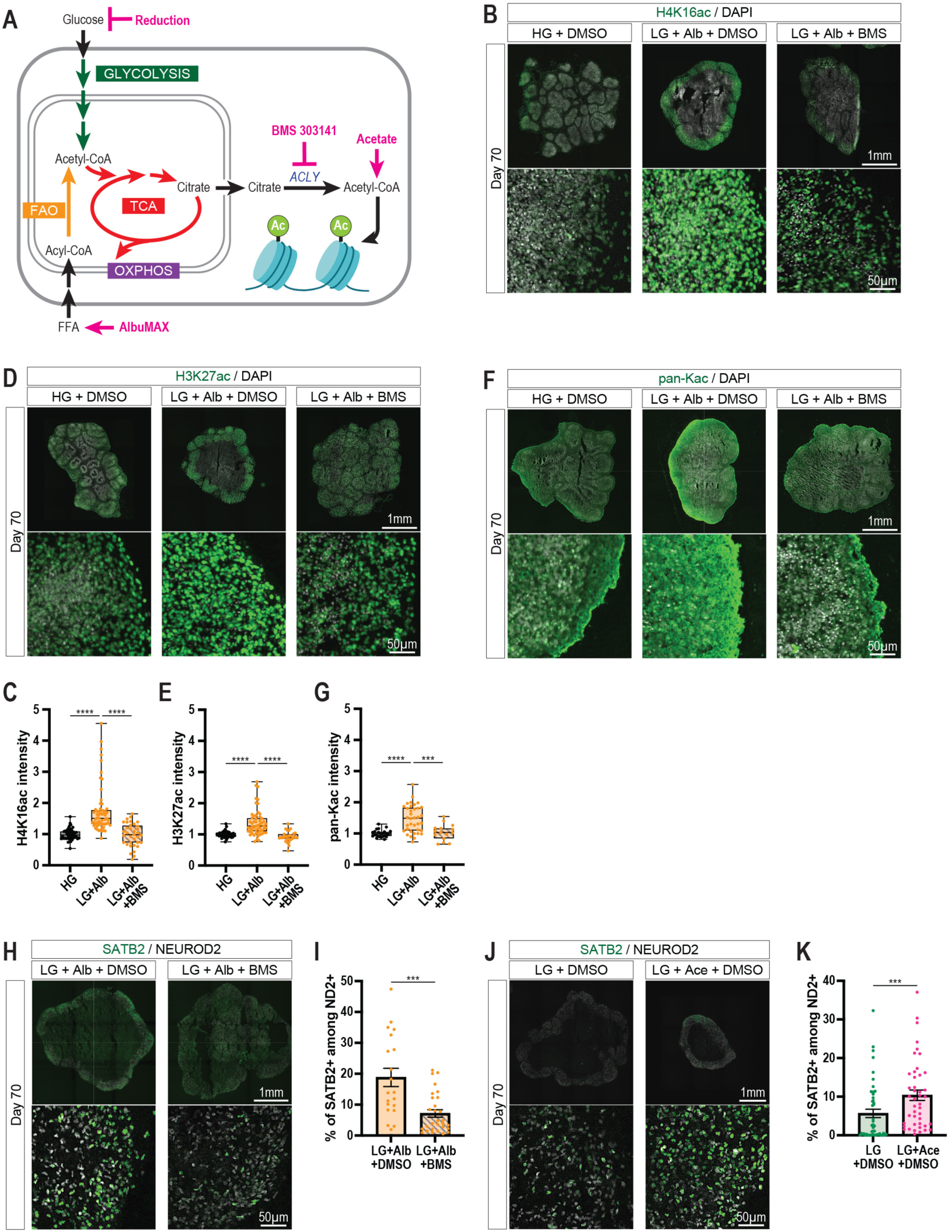
FAO accelerates temporal patterning through protein acetylation dynamics. (A) Schematic of metabolic pathways targeted by indicated chemical compounds. ATP citrate lyase (ACLY). (B) Representative images of H4K16 acetylation in cortical organoid at day 70. (C) Quantification of mean signal intensity of H4K16ac from each organoid. Each data point represents an individual organoid. High glucose with DMSO (HG+DMSO): N=44 organoids from eight independent differentiation batches. Low glucose with 0.5% AlbuMAX and DMSO (LG+Alb+DMSO): N=60 organoids from eight independent differentiation batches. Low glucose with 0.5% AlbuMAX and 5µM BMS (LG+Alb+BMS 303141: N=35 organoids from eight independent differentiation batches. Dunn’s multiple comparisons test. (D) Representative images of H3K27 acetylation in cortical organoid at day 70. H4K16 acetylation. (E) Quantification of mean signal intensity of H3K27ac from each organoid. Each data point represents an individual organoid. High glucose with DMSO (HG+DMSO: N=36 organoids). Low glucose with 0.5% AlbuMAX and DMSO (LG+Alb+DMSO: N=55 organoids from seven independent differentiation batches. Low glucose with 0.5% AlbuMAX and 5µM BMS (LG+Alb+BMS 303141: N=28 organoids from seven independent differentiation batches. Dunn’s multiple comparisons test. (F) Representative images of pan-K acetylation immunostainings in cortical organoids at day 70. (G) Quantification of mean signal intensity of pan-K ac from each organoid. Each data point represents an individual organoid. High glucose with DMSO (HG+DMSO: N=23 organoids from six independent differentiation batches. Low glucose with 0.5% AlbuMAX and DMSO (LG+Alb+DMSO): N=36 organoids from six independent differentiation batches. Low glucose with 0.5% AlbuMAX and 5µM BMS (LG+Alb+BMS 303141): N=15 organoids from six independent differentiation batches. Dunnett’s multiple comparisons test. (H, J) Representative images of cortical organoids at day 70. (I) Quantification of the proportion of SATB2-positive cells. Each data point represents an individual organoid. Low glucose with 0.5% AlbuMAX and DMSO (LG+Alb): N=20 organoids. Low glucose with 0.5% AlbuMAX and 5µM BMS (LG+Alb+BMS 303141): N=29 organoids from three independent differentiation batches. Mann-Whitney test. (K) Quantification of the proportion of SATB2-positive cells. Each data point represents an individual organoid. Low glucose with DMSO (LG+DMSO): N=48 organoids from five independent differentiation batches. Low glucose with Acetate (100 µM) and DMSO (LG+Ace+DMSO): N=44 organoids from five independent differentiation batches. Mann-Whitney test. *P < 0.05, **P < 0.01, ***P < 0.001, ****P < 0.0001.

To test whether this model could apply to cortical temporal patterning we first measured Ac-CoA levels in mouse and human cortical cells prepared as above from mouse embryonic brain and human cortical organoids (Fig. 4F). Remarkably this revealed that the levels of Acetyl-CoA were lower in human than mouse cortical cells (Fig. 4G), in line with the lower FAO activity (Fig. 4, B and E). Next we quantified Acetyl-CoA in cortical organoids cultured in control HG conditions of with Alb treatment as above (Fig. 4H). This revealed that the levels of Acetyl-CoA were higher in the Alb conditions (Fig. 4I), thus strongly pointing towards Acetyl-CoA as a main metabolite increased in response to FAO during corticogenesis.

We next examined the impact of Alb treatments on protein acetylation patterns in cortical organoids. This revealed an increase in histone H3 lysine 27 (H3K27) and histone H4 lysine 16 (H4K16) acetylation marks (Fig. 5, B to E). Notably we also observed increased levels of protein acetylation in the FFA-treated condition (Fig. 5, F and G), indicating that other proteins than histones are acetylated following FFA treatments. Importantly the FFA-induced increase of acetylated proteins were all blocked by simultaneous treatment with ACLY inhibitors (BMS-303141: (referred to as BMS), that prevents the conversion of citrate into Acetyl-CoA (Fig. 5, B to G). Thus, FFA treatment in cortical organoids leads to increased protein acetylation dynamics through increased citrate to acetyl-CoA conversion.

Finally test whether the molecular effects of FAO on acetylation were relevant to the acceleration of cortical temporal patterning, we examined temporal neuronal fate markers in cortical organoids treated with Alb alone or together with BMS. Remarkably, BMS blocked all the effects of FFA on accelerated temporal patterning, indicating that the conversion of citrate into Acetyl-CoA is required for the effects of FAO on temporal patterning (Fig. 5. H and I, and fig. S8. A and B). Moreover to test whether acetylation dynamics was sufficient to accelerate temporal patterning, we treated cortical organoids with exogenous acetate, which can constitute a source of Acetyl-CoA bypassing FAO metabolism (*52*). Acetate treatment also led to precocious generation of late fate SATB2-positive neurons, although at lower levels than FAO, indicating that other metabolites may link FAO and temporal patterning (Fig. 5. J and K, and fig. S8. C and D).

Overall, these data indicate that mitochondria FAO metabolic rates set the timescale of temporal patterning at least in part through increased TCA-dependent Ac-CoA generation that leads to enhanced protein acetylation including histone marks of transcriptional regulation.

## Discussion

Here we found that mitochondrial fatty acid oxidation (FAO) rates are higher in mouse than human cortical neural progenitor/stem cells, in which they constitute a robust global accelerator of human corticogenesis, at both cellular and molecular levels. FAO was found to act through Acetyl-CoA-dependent enhanced protein acetylation dynamics, including on specific Histone marks associated with transcriptional regulation. Our data thus indicate that the scale of temporal patterning of corticogenesis, which is considerably prolonged in the human species, is set in part by species-specific rates of mitochondrial fatty acid metabolism upstream of protein acetylation and epigenetic remodeling.

Mitochondria metabolism was previously found to be essential for the balance of self-renewal and differentiation of neural stem cells, form fly to human species, mostly in link with oxidative phosphorylation-generated and reactive oxygen species and NAD+ (*16*, *23*, *53*). Here we find that mitochondrial metabolism influences developmental timing and temporal patterning through TCA-linked Acetyl-CoA-dependent protein acetylation rates, rather than through the oxidative phosphorylation. Our data are in line with the observation that in the presomitic mesoderm oscillations, which are also regulated by species-specific metabolic rates (*15*), the rates of ATP production *per se* do not appear to be correlated with the tempo of the somitic clock across multiple species (*11*). It will be interesting to determine to which extent TCA metabolites and protein acetylation rates also contribute to set the tempo in other systems, in parallel or together with other proposed mechanisms such as protein turnover (*15*, *54*).

Our data identify FAO as the most efficient metabolic pathway to accelerate corticogenesis. This is in line with the highly efficient capacity of FAO to enhance Acetyl-CoA generation, protein acetylation, and thereby to regulate cell fate transitions (*47*, *48*, *52*, *55–57*). Indeed, our data show that enhanced FAO during human corticogenesis leads to upregulated levels of Acetyl-CoA and acetylation of proteins. These could include non-histone proteins such as specific transcriptional or cell regulators (*47*, *57–61*). As for histones, we found higher levels of H4K16 and H3K27 acetylation marks, which correspond typically to active enhancer/promoter regions. Our data thus point to epigenetic chromatin remodeling as a key substrate for the global transcriptomic changes following enhanced FAO in corticogenesis. Our data point to a model whereby metabolism-dependent-turnover of acetylation of histones could accelerate the temporal progression of the developmental GRNs. On the other hand, acetylation may not be the only histone post-translational modification involved the species-specific timescale of corticogenesis. The speed of maturation of human cortical post-mitotic neurons was recently shown to be slowed down by an epigenetic brake, relying on histone methylation rates regulated by the Polycomb complex, although it remains unclear how much this block is species-specific (*62*). The Polycomb complex is also required for neural temporal patterning in the fly and the mouse, acting to prevent the transition from early to late neuronal fates (*63*). In this context it will be interesting to determine if FAO could influence species-specific developmental tempo through histone methylation as well, as it can also be regulated by TCA-derived metabolites (*49*, *64*), or whether histone acetylation and methylation dynamics influence different aspects of temporal patterning.

Importantly, we also observed temporal effects of low glucose treatment on precocious expression of late neuronal fate markers, although at much lower levels than those observed with FFA treatment, as well as LDHA inhibition on precocious gliogenesis. Thus, while our data suggest that FAO seems to be the main source of metabolites that influence temporal patterning, other metabolic pathways or products could be involved as well, such as pentose phosphate metabolism and lactate signaling, which have been previously shown to modulate mouse and human cortical neurogenesis (*19*, *27*, *28*).

The sequential generation of cortical neurons of distinct identities is highly conserved between species, while the timescale of this temporal pattern differs between species. Consistent with this notion, our data support a mechanism by which metabolic and epigenetic rates set the scale of temporal patterning but without altering the structure of the GRN underlying neuronal specification, which is likely to be largely conserved between species. On the other hand, human- specific changes were previously shown to impact the tempo of corticogenesis, including cis and trans regulatory elements as well as Notch and APP signaling pathways (*65–69*). It will be interesting to test in the future whether and how any of these species-specific changes can be linked to the lower metabolic rates that we uncovered here. It will be also important to relate the metabolic pathways uncovered here to outer-radial glial cells, key contributors to the expansion of the human cortex (*43*), which express markers that we found to be upregulate by increased FAO. Along the same lines, the upstream mechanisms that underlie the species differences in metabolism of cortical RGCs remain unknown. These could include protracted and/or lower expression of mitochondrial genes, as previously observed for neuronal maturation (*13*), but also likely involve the numerous levels of post-transcriptional regulation of metabolic pathways. Species-specific patterns of metabolism in RGCs could be also linked to recently described regional differences in expression of mitochondrial genes in the mouse embryonic brain (*18*).

Finally, our data illustrate the importance of metabolism as a critical parameter to optimize in vitro systems of modeling of development, which are likely to be influenced by in vivo vs in vitro context (*70*). They also open novel opportunities to accelerate the tempo of human neural organoid development, which could be of great interest for the modeling of neural diseases or the induction of clinically relevant cell fates such as astrocytes, which have been hindered by the prolonged timescale of human brain development.

## Acknowledgements

We thank the PV lab members for their helpful discussion and invaluable assistance. We thank the VIB CBD Single Cell Expertise Unit and the VIB Single Cell Core for snRNA-seq experiments and the VIB BioImaging Core for image acquisitions and automated analyses. We thank Ekaterina Epifanova and Anne Firquet for assistance with collection of human fetal tissue.

## Author contributions

Conceptualization and Methodology, RI, IGL, EE, NP, SA, SMF, PV; Investigation, RI, IGL, EE, RL, MD, VL, AV, MP, SMF, PV.; Formal Analysis, RI, IGL, EE, NC, SP, PVM, BG, KD, BG, SMF, SC, PV; Writing, RI, IGL, EE, PV; Funding acquisition, PV; Resources, PVe, AB, LN, PV; Supervision, PV.

## Materials and Methods

### Mice

All mouse experiments were performed with the approval of the KU Leuven Committee for animal welfare. Mice were housed under standard conditions (12h light:12h dark cycles) with food and water ad libitum. Embryos (aged E12.5 - E16.5) of the mouse strain ICR (CD1, Charles River Laboratory) or Swiss (Janvier Labs) were used for in utero experiments and primary cultures. The plug date was defined as embryonic day E0.5, and the day of birth was defined as P0. The data obtained from all embryos were pooled without discrimination of sexes. RAG2 knock-out mice were used for xenotransplantation experiments.

### Lentiviral preparation

HEK293T cells were transfected with the packaging plasmids, psPAX2, a gift from Didier Trono (Addgene plasmid # 12260; http://n2t.net/addgene:12260; RRID:Addgene_12260) and pMD2.G, a gift from Didier Trono (Addgene plasmid # 12259; http://n2t.net/addgene:12259; RRID:Addgene_12259), and a plasmid of the gene of interest in the lentiviral backbone pLenti-human Synapsin I promoter (hSynI)-EmGFP-WPRE (*1*). Three days after transfection, the culture medium was collected and viral particles were enriched by filtering (Amicon Ultra-15 Centrifuge Filters, Merck, Cat#UFC910008). Titer check was performed on HEK293T cell culture for every batch of lentiviral preparation

### Cell culture media

hES medium (Wisconsin medium):

This medium was used to maintain and expand human PSC.

KnockOut-DMEM (Thermo Fisher Scientific, Cat#10829018) with KnockOut Serum Replacement (20%, Thermo Fisher Scientific, Cat#10828028), Non-Essential Amino Acids Solution (1x, Thermo Fisher Scientific, Cat#11140050), L-Glutamine (2mM, Thermo Fisher Scientific, Cat#25030081), 2-Mercaptoethanol (100µM, Merck, Cat#M3148) and Human FGF-basic (10ng/ml, PeproTech, Cat#AF-100-18B).

DDM/B27 medium:

DMEM/F12 + GlutaMAX (Thermo Fisher Scientific, Cat#10565042) with N2-supplement (1x, Thermo Fisher Scientific, Cat#A1370701), B27 supplement minus Vitamin A (1x, Thermo Fisher Scientific, Cat#12587010), Bovine Albumin Fraction V (0.05%, Thermo Fisher Scientific, Cat#15260037), 2-Mercaptoethanol (100µM, Merck, Cat#M3148), Non-Essential Amino Acids Solution (1x, Thermo Fisher Scientific, Cat#11140050) and Sodium Pyruvate (1mM, Thermo Fisher Scientific, Cat#11360070).

Nb/N2B27 high glucose (25 mM) medium:

This medium was used for mouse primary culture and human cortical organoid culture from day 30.

Neurobasal-A (Thermo Fisher Scientific, Cat#A2477501) with B27 supplement (1x, Thermo Fisher Scientific, Cat#17504044), N2 supplement (1x Thermo Fisher Scientific, Cat#117502048), Sodium Pyruvate (0.227 mM, Thermo Fisher Scientific, Cat#11360070), Penicillin-Streptomycin (50 U/ml, Cat#1570063), Glucose (25 mM, Thermo Fisher Scientific, Cat#A2494001) and GlutaMAX supplement (1x, Thermo Fisher Scientific, Cat# 35050061).

Nb/N2B27 low glucose (2.5 mM) medium:

This medium was used for human cortical organoid culture from day 30.

Neurobasal-A (Thermo Fisher Scientific, Cat#A2477501) with B27 supplement (1x, Thermo Fisher Scientific, Cat#17504044), N2 supplement (1x Thermo Fisher Scientific,

Cat#117502048), Sodium Pyruvate (0.227 mM, Thermo Fisher Scientific, Cat#11360070), Penicillin-Streptomycin (50 U/ml, Cat#1570063), Glucose (2.5 mM, Thermo Fisher Scientific, Cat#A2494001) and GlutaMAX supplement (1x, Thermo Fisher Scientific, Cat# 35050061).

### Human PSC culture and cortical organoid differentiation

Human embryonic stem cells (hESC) (H9; WiCell, Cat #NIHhESC-10-0062; female donor) were maintained in mTeSR1 medium (Stem Cell Technologies, Cat #85850) on growth factor-reduced Matrigel substrate (Corning, Cat #354230) in a feeder-free condition. hESC were passaged as clumps using ReLeSR (Stem Cell Technologies, Cat #100-0483) once or twice per week and used for cortical organoid differentiation within eight passages after thawing.

Human induced pluripotent stem cells (hiPSC), AH4 line (female donor), were maintained on irradiated MEFs in Wisconsin medium. Cells were passaged enzymatically as clumps with Collagenase type IV (1 mg/ml, Thermo Fisher, Cat #17104019) and Dispase II (1 mg/ml, Thermo Fisher, Cat #17105041) every three to five days and used for cortical organoid differentiation within five passages after thawing.

Both hESC and hiPSC lines were regularly checked for mycoplasma by PCR and frozen in mFreSR medium (Stem Cell Technologies, Cat #05855) in an isopropanol freezing container at -80°C overnight and subsequently in a liquid nitrogen tank, H9 hESC-derived cortical organoids were generated based on previously (Cederquist et al., 2019) with slight modification. Two days before starting neuronal cell differentiation (day -2), hESC were dissociated to single cells using Accutase (Thermo Fisher, Cat #A1110501), seeded at 12,000 cells in 100 µl per well in V-bottom cell-repellent microwells (Greiner Bio-One, Cat #651970) and allowed to aggregate during two days in mTeSR1 medium in the presence of 10µM ROCK inhibitor (Merck, Cat #688000) at 37°C and 5% CO_2_. On day 0 of differentiation, medium was replaced with Essential 6 medium (Thermo Fisher, Cat #A15165-01) containing 100 nM LDN193189 (Stem Cell Technologies, Cat #72147), 10 µM SB431542 (Stem Cell Technologies, Cat #72234) and 5 µM XAV939 (Stem Cell Technologies, Cat #72674). Medium was renewed every other day. At day 8, spheroid bodies were transferred into droplets containing of 100 µl growth factor-reduced Matrigel and 67 µl DDM/B27 medium, allowed to gelify during 1 hr in well centers of cell-repellent 6-well plates (Greiner Bio-One, Cat #657970) and further supplied DDM/B27 medium with Penicillin-Streptomycin (50 U/ml, Cat#1570063) and Insulin (2.5 mg/ml, Sigma-Aldrich, Cat #I9278). At day 12 or 13, organoids were mechanically released from the droplets by resuspension pipetting with a serological 5 ml-pipet and maintained in the same culture vessels on an orbital shaker (100 rpm) in an identical configuration, except containing B27 (Thermo Fisher, Cat #17504044) instead of B27 without vitamin A, at 37°C and 5% CO_2_. Medium was renewed completely three times per week.

AH4 iPSC-derived cortical organoids were generated as described in van Benthem et al BioRxiv (doi: https://doi.org/10.1101/2023.09.25.559314). Entire hiPSC colonies were detached from the MEF feeder layer, washed in Wisconsin medium without bFGF and placed in cell-repellent 6-well plates in complete neural induction medium, DMEM/F12 + GlutaMAX with 20% KnockOut Serum Replacement, 1% MEM-NEAA, 50 U/ml Penicillin-Streptomycin, 100 µM 2-Mercaptoethanol, 100 ng/ml Noggin (Bio-techne, Cat #1967-NG), 10 µM SB431542, 2 µM XAV939 and 10 µM Y-27632. Neural induction medium without Y-27632 was renewed on day 1 and every other day onwards until day 16. At day 17, embryoid bodies were gently collected in 5 ml differentiation medium, DMEM/F12 + GlutaMAX supplemented with 1% MEM-NEAA, 50 U/ml Penicillin-Streptomycin, 100 µM 2-Mercaptoethanol, 2.5 µg/ml insulin, 1% N2 supplement, and 2% B27 without vitamin A, placed in cell-repellent 6-well plates and maintained on an orbital shaker (100 rpm) at 37°C and 5% CO_2_. Medium was renewed completely three times per week.

At day 30, medium was switched to Nb/N2B27 high or low glucose (25 mM or 2.5 mM) medium. From day 45 onwards, 1% vol/vol growth factor-reduced Matrigel was added to the culture medium. Medium was renewed completely three times per week.

### Chemical compound treatment

From day 30, the following compounds at the specified concentrations were added to the Nb/N2B27 medium. AlbuMAX I (0.5% w/vol, Thermo Fisher, Cat #11020021). Etomoxir (50 µM, Merck, Cat #E1905). UK-5099 (1.25 µM, Merck, Cat #5048170001). BMS-303141 (5µM, TOCRIS, Cat #4609). GSK-2837808A (5 µM, MedChemExpress, Cat #HY-100681).

Sodium acetate (100 µM, Merck, Cat #HY-100681). Palmitic acid (100µM, Merck, Cat #P0500) with fatty-acid free Albumin (Merck. Cat #126575). Dimethyl sulfoxide (DMSO, Merck, Cat #D2650) concentration was maintained at 0.025% vol/vol. Medium was renewed completely three times per week.

For 2D culture, from day 25, cells were treated with 0.5% AlbuMAX I and 5µM GSK-2837808A in Nb/N2B27 low glucose medium or 0.025% DMSO in Nb/N2B27 high glucose medium until the day of oxygraphy measurement.

### Human PSC culture and 2D cortical cell differentiation

Human ESC (hESC) (H9; WiCell Cat # NIHhESC-10-0062; female donor) were grown on irradiated mouse embryonic fibroblasts in the ES cell medium until the start of cortical cell differentiation. Cortical cell differentiation was performed as described previously (*1*). Two days before starting neuronal cell differentiation (day -2), hESCs were dissociated with Accutase (Thermo Fisher Scientific, Cat#00-4555-56) and plate on Matrigel-coated (BD, Cat#354277) plates at low confluency (5,000 cells/cm^2^) in hES medium with 10µM ROCK inhibitor (Merck, Cat#688000). On day 0 of the differentiation, the medium was changed to DDM/B27 medium with recombinant mouse Noggin (100 ng/ml, R&D systems, Cat#1967-NG). The medium was changed every other day until day 6. From day 6, the medium was changed every day until day 16. At day 16, medium was changed to DDM/B27 medium and changed every day until day 25. At day 25, the differentiated cortical cells were dissociated using Accutase and cryopreserved in mFreSR (STEMCELL technologies, Cat#05855). Differentiated cortical cells were validated for neuronal and cortical markers by immunostaining using antibodies for TUBB3 (1:2,000; BioLegend, Cat #MMS-435P), TBR1 (1:1,000; Abcam, Cat #ab183032), CTIP2 (1:1,000; Abcam, Cat #ab18465), FOXG1 (1:1,000; Takara, Cat #M227), SOX2 (1:2,000; Santa Cruz, Cat #sc-17320), FOXP2 (1:500; Abcam, Cat #ab16046), SATB2 (1:2,000; Abcam, Cat #ab34735).

### Mouse primary cortical cell culture

Timed pregnant mouse embryos at E12.5-E16.5 were used. Cerebral cortices were dissected and enzymatically dissociated using Trypsin-EDTA (Thermo Fisher, Cat #25300054) with DNase I (VWR, Cat #ROCKMB-101-0100). The tissue was triturated with a glass Pasteur pipette to generate a single-cell suspension. The single cells were plated on laminin- and poly-D-lysine-coated coverslips or cell culture well plate (12-well or 6-well) at high confluency (400,000 to 500,000 cells/cm^2^) using Nb/N2B27 high glucose (25 mM) medium.

### Human fetal cortical cell culture

Human fetal cortex was obtained after dissection to remove skin and skull. The samples were preserved in pre-oxygenated artificial cerebrospinal fluid, and no more than 3.5 hrs elapsed from extraction to dissociation. Gestational age ranged from week 10 to 13.

For dissociation, the tissue was resuspended in 500 µl of Hank’s Balanced Salt Solution (HBSS) and Liberase DH (Roche, Cat #5401054001) at 0.28 U/ml working concentration and incubated for 40 min at 37°C on a shaking platform. To achieve complete tissue dissociation, a second reaction was performed using the Papain Dissociation System (Worthington Biochemical Corporation, Cat #LK003150) following the manufacturer’s protocol with minor variations. Papain (20 U/ml) and DNase I (2000 U/ml) were added to the previous reaction and incubated for 20 min at 37°C on a shaking platform. The tissue was mechanically dissociated with a glass Pasteur pipette until a uniform cell suspension was achieved. Small tissue clumps were removed. After centrifugation at 300 xg for 5 minutes, cells were resuspended in DNase I-diluted albumin-inhibitor solution and layered on 5 ml of Ovomucoid solution to completely stop Papain activity. Following a final centrifugation at 300 xg for 5 minutes, cells were resuspended in Nb/N2B27 high glucose medium and plated on laminin-, poly-D-lysine-, and Matrigel-coated 6-well plates at high confluency (400,000-500,000 cells/cm²). Cells were cultured for 2 days before undergoing oxygraph.

### Immunostaining

Organoids were transferred into a 15 ml-tube, washed once in PBS at room temperature and fixed in 4% paraformaldehyde (PFA) in PBS overnight at 4°C. The following day, organoids were washed three times with PBS on gentle agitation and then transferred into 30% sucrose PBS overnight at 4°C for cryoprotection. The next day, organoids were embedded in Tissue Freezing Medium (Leica, Cat #14020108926) and frozen using dry ice. Tissue blocks were stored at -80°C until sectioning. Cryosectioning was performed using a cryostat (Leica 3050S) in 10 or 20 µm thickness under RNase-free conditions and mounted onto Superfrost slides (Thermo Fisher, Cat #J1800AMNZ). Cryosections were washed during 10 min prior to a 1 hr permeabilization and blocking step in PBS containing 3% BSA, 1% horse serum and 0.3% Triton X-100, and subsequent primary antibodies incubation in the same solution overnight at 4°C. The sections were immunostained with following antibodies: NEUROD2 (1:500 , Abcam, Cat #ab104430), TBR2 (1:1,000, Abcam, Cat #ab216870), PAX6 (1:500, BD Biosciences, Cat #561-462), SOX2 (1:200, R&D systems, Cat #AF2018), TBR1 (1:200, Abcam, Cat#ab31940), CTIP2 (1:500, Abcam, Cat#ab18465), SATB2 (1:50, Abcam, Cat#ab51502), H3K27ac (1:500, Thermo Fisher, Cat #39133), H4K16ac (1:500, Abcam, Cat #ab109463), and pan-Kac (1:500, PTM Bio, Cat #PTM-101).The following day, cryosections were washed three times with PBS, incubated at room temperature for 3 hrs with appropriate secondary antibodies and DAPI (Sigma, Cat #D9542), washed again three times with PBS and mounted with a coverglass using Glycergel mounting medium (Dako, Cat #C0563).

For immunocytochemistry of coverslips, cells were fixed with 4% PFA in PBS 1h at 4°C. The coverslips were transferred into PBS, then blocked using PBS with 3% horse serum (Thermo Fisher, Cat #16050122) and 0.3% Triton X-100 (Sigma-Aldrich, Cat #T9284) during 1 h, and incubated overnight at 4°C with the primary antibodies. After three washes with PBS/0.1% Triton X-100, the coverslips were incubated in PBS for 5 min at room temperature and incubated 2 h at room temperature with the appropriate secondary antibodies with DAPI (Merck, Cat #D9542). The coverslips were again washed three times with PBS for 5 min. The coverslips were mounted on a Superfrost slide (Thermo Fisher) using Glycergel mounting medium (Dako, Cat #C0563). The cells were immunostained with the following antibodies for GFP (1:2,000, Abcam, Cat #ab13970), NEUROD2 (1:1,000, Abcam, Cat #ab104430), TUBB3 (1:2,000, BioLegend, Cat #MMS-435P), TUBB3 (1:2,000, BioLegend, Cat #PRB-435P), TBR2 (1:1,000, Abcam, Cat #ab216870), PAX6 (1:1,000, BD Biosciences, Cat #561-462), SOX2 (1:200, R&D systems, Cat #AF2018), TBR1 (1:1,000, Abcam, Cat#ab31940).

### Xenotransplantation of cortical organoid-derived human neurons

H9 hESC were transduced with LV-hSynI-EmGFP-WPRE at the moment of cell seeding (Day -2) in mTeSR 1 medium. On day 0, the lentivirus-containing medium was replaced with Essential 6 medium (Thermo Fisher, Cat #A15165-01) containing 100 nM LDN193189, 10 µM SB431542 and 5 µM XAV939. Further culturing steps were done as described above (Section: Human PSC culture and cortical organoid differentiation). One week before the xenotransplantation day, organoids were collected and washed in oxygenated HBSS (95% O2, 5% CO2), then snapped into two pieces with a sterile blade to initially remove debris from the organoid cores. Organoids were subsequently washed twice more in HBSS and incubated for 30 minutes at 37°C in pre-warmed oxygenated Papain / DNase I (Worthington Biochemical Corporation, Cat #LK003150) on an orbital shaker (100 rpm). A single cell suspension was prepared by gentle trituration and centrifugation at 300 xg for 5 min. Debris was removed by subsequent gradient centrifugation of Ovomucoid protease inhibitor / DNase I at 70 xg for 6 min. The cells were resuspended in Nb/N2B27 medium. After cell counting and viability assessment with Trypan blue, live cells were seeded at high density (600,000 to 700,000 cells/cm²) in growth factor-reduced Matrigel-coated 6-well plates. Medium was renewed every other day. At 3 days before xenotransplantation, cells were treated with 10 µM gamma-secretase inhibitor (DAPT, Abcam, Cat #ab120633) and cultured three additional days. On the xenotransplantation day, the cortical cells were dissociated using NeuroCult Enzymatic Dissociation Kit (STEMCELL technologies, Cat #05715) following manufacturer’s instructions and suspended in the injection solution containing 20mM EGTA (Merck, Cat #03777) and 0.1% Fast Green (Merck, Cat #210-M) in PBS at 100,000–200,000 cells/µl. Approximately 1-2 µl of cell suspension was injected into the lateral ventricles of each hemisphere of neonatal (postnatal day 0 or 1) immunodeficient mice (Rag2 knockout) using glass capillaries pulled on a horizontal puller (Sutter P-97).

At 14 days post-transplantation, mice were perfused with 4% PFA and 0.5% GA in 0.1M phosphate buffer, and the brains were collected. Brains were dissected, and 100 μm sections were prepared using a Leica VT1000S vibratome. Slices were transferred into PBS with sodium azide (0.5 μg/ml: Sigma-Aldrich, Cat #71289), then blocked with PBS with 3% horse serum and 0.3% Triton X-100 during 1 hr, and incubated three overnight at 4°C with the primary antibodies. After three washes with PBS/0.1% Triton X-100, slices were incubated in PBS for 1 hr at room temperature and incubated 2 hrs at room temperature with the appropriated secondary antibodies with DAPI. Sections were again washed three times with PBS for 1 hr. The sections were mounted on a Superfrost slide and then added Glycergel mounting medium. The slices were immunostained using antibodies for GFP (1:2,000; Abcam, Cat #ab13970), SATB2 (1:2,000; Abcam, Cat #ab34735), and CTIP2 (1:1,000; Abcam, Cat#ab18465). Imaging of xenotransplanted tissues (Fig. 2K) was performed using a Zeiss LSM880 confocal microscope with a 10x objective.

### Image acquisition

Organoid sections were imaged using a Nikon Ti2 inverted microscope equipped with a Cicero spinning-disk confocal module (CrestOptics S.p.A.) and a Kinetix sCMOS camera (Teledyne Photometrics). The system was operated via NIS-Elements software (versions 6.10.0 and 6.10.01, Nikon Instruments Europe B.V.).

Excitation was provided by a Celesta light engine (Lumencor) and emission was collected using a Penta filter set with bandpass filters. A 20× Plan Apochromat Lambda D air objective (NA 0.8) was used for all acquisitions.

### Image processing and quantification

Images were stitched using NIS-Elements software (versions 6.10.0 and 6.10.01, Nikon Instruments Europe B.V.). Processing and quantification were performed using a combination of FIJI (ImageJ 1.54p with Java 1.8.0_322, 64-bit; https://fiji.sc/) and the ijp-kheops, along with QuPath (version 0.5.1), following a semi-automated pipeline for nuclear segmentation and marker quantification.

Briefly, raw multichannel .nd2 images were processed in FIJI using a custom macro script. This script enhanced the channel of interest to facilitate nuclear segmentation by applying a rolling ball background subtraction (radius: 50 pixels). The enhanced channel was appended as a fifth channel to the original image stack. The resulting images were saved in pyramidal OME-TIFF format for downstream analysis.

Processed images were imported into QuPath as part of a project. Regions of interest (ROIs) were manually drawn for each image to select the entire surface of the organoid, avoiding the center that contains mostly necrotic cells. A custom Groovy script was used for nuclear segmentation and marker quantification. Nuclear segmentation was performed using StarDist (https://github.com/stardist) via the QuPath StarDist extension (https://github.com/qupath/qupath-extension-stardist), based on the enhanced channel. Segmented nuclei were filtered by area and intensity thresholds and converted into circular ROIs centered on the nuclear centroid (*74,75*).

Mean fluorescence intensities were measured across all channels within these ROIs. A composite classifier was applied to categorize cells based on marker expression.

Quantification results were exported from QuPath as tab-separated values (TSV) files for further analysis.

For full implementation details and reproducibility, the complete pipeline and scripts are available at the associated GitHub repository: https://github.com/vib-bic-projects/202504_Isabel_NuclearClassification

### Nuclei extraction and snRNA sequencing

The nuclei for 10x Genomics Chromium GEM-X Single Cell 3’ v4 (CG000731 | Rev B) were extracted from human organoid samples that were snap-frozen in liquid nitrogen and stored at -80 °C. We transferred the frozen organoids to 500 μl of ice-cold homogenization buffer (10 mM Tris-HCl pH 7.5, 150 mM NaCl, 22 mM MgCl_2_, 1 mM CaCl_2_, 250 mM sucrose, 25 mM KCl, 0.5% BSA, 0.03% tween-20, 1× complete protease inhibitor, 1 mM DTT and 1 U/μl Rnase in plus) in a Dounce Homogenizer (KIMBLE, 1 ml). The samples were left to thaw on ice for 3 min before being homogenized. The samples were incubated for 5 min on ice, during which time the homogenate was filtered through a 70 μm cell strainer (Corning). The filtrate was transferred to a 1.5 ml DNA LoBind tube (Eppendorf) and centrifuged at 500 xg for 5 minutes. The supernatant was discarded, and the pellet was resuspended with 520 μl with wash buffer 1 (10 mM Tris-HCl pH 7.5, 150 mM NaCl, 22 mM MgCl_2_, 1 mM CaCl_2_, 250 mM sucrose, 25 mM KCl, 0.5% BSA, 1× complete protease inhibitor,1 mM DTT and 1 U/ μl Rnase in plus). Next, we added 520 μl gradient medium (50% Optiprep, 10 mM Tris-HCl pH 7.5, 1 mM CaCl_2_, 5 mM MgCl_2_, 75 mM sucrose, 1× complete protease inhibitor, 1 mM DTT and 1 U/μl Rnase in plus) and mixed the entire volume by gentle pipetting. In a 2 ml DNA LoBind tube (Eppendorf) we layered 790 μl Optiprep cushion (29% Optiprep, 30 mM Tris-HCl pH 8, 75 mM KCl, 15 mM MgCl_2_, 125 mM sucrose) at the bottom and 1040 μl of the sample on top, gently without disrupting the cushion. The tube was then centrifuged at 9,000 xg for 20 minutes at 4 °C. After the centrifugation the debris on top was carefully removed with a 1 ml tip and the rest of the supernatant was also carefully removed. The pellet was gently resuspended in 80 ul of resuspension buffer (PBS with 2% BSA and 1 U/μl Rnase in plus). Nuclei quality and concentration were assessed by the LUNA-FL Dual Fluorescence Cell Counter. 20,000 nuclei were loaded for the RT reaction for each sample according to the 10x Genomics Chromium GEM-X Single Cell 3’ v4 (CG000731 | Rev B).

### Pre-processing of snRNAseq data

FASTQ files of snRNA-seq libraries were mapped to refdata-GRCh38-2024-A index with 10x Genomics Cell Ranger v9.0.0 *count* command.

### SnRNAseq analysis of cortical organoids

Filtered gene expression matrices were processed using the Scanpy (1.9.8) package (*69*). Nuclei with less than 500 genes or more than 10% of counts present in mitochondrial genes were excluded, counts were log-normalized and counts per nuclei were scaled to a total of 10,000. Top 1500 highly variable genes were selected with *pp.highly_variable_genes* using seurat_v3 flavor and principal component analysis (PCA) was performed with the selected genes. PCs were adjusted between *treated* and *untreated* data using Harmony (*70*) with *external.pp.harmony_integrate.* Top 15 adjusted PCs were then used to compute a neighborhood graph with n_neighbours=15. UMAP was computed using default Scanpy parameters. Clusters were annotated based on known marker genes (*36–38*).

RGC: PAX6, SOX2,HES1,HOPX, IPC: EOMES, NEUROD4, Young Neurons: NEUROD2, NEUROD6, SOX4, SOX11, Non-IT immature: NFIA, TBR1, SOX4, NEUROD2, BCL11B, IT-immature: SOX4, NEUROD2, SATB2, CUX2, IT ExN: MEF2C, SATB2, CUX2, GRIN2B, CAMK2B, Non-IT ExN: TLE4, BCL11B, SOX5, MEF2C, GRIN2B, CAMK2B, IN: GAD2, GAD1, C. plexus: TTR.

Late RGC (499 genes) and early RGC (204 genes) gene scores were obtained from (*41*). Gene scores were calculated using *tl.score_genes* in which the number of control genes were always set to the size of the gene list used for scoring. Unsupervised clustering was performed using the Leiden algorithm at default resolution on integrated data. Differential expression tests were performed between corresponding organoid age and cell types using PyDESeq2 with 3 pseudobulks per condition (*71*). Gene ontology (GO) analyses were performed on differentially expressed genes (higher expression: Log2FC>0.25, FDR<0.01, lower expression: Log2FC<-0.25, FDR<0.01) using GSEApy (1.1.3) package on GO Biological Process 2025 gene set using the *enrichr* function (*72*).

### Integration of organoids and human developing cortex cells

The data of developing human cortex (*10*) was downloaded from NeMO archive (https://assets.nemoarchive.org/dat-oiif74w). The expression matrix was subset to select only excitatory neurons and progenitors and stages from first trimester to third trimester cells. Organoids and *in vivo* expression matrices were concatenated and log-normalized. Top 1500 Highly variable genes were selected for PCA. PCs were adjusted between Organoids and *in vivo* data using Harmony with *external.pp.harmony_integrate*. Corrected PCA embeddings were then used to form the neighborhood graph and UMAP was calculated with the default Scanpy parameters.

### Label transfer of human in vivo stages to organoid cells

In order to predict stages of organoid RGCs, a SCANVI (Xu et al., 2021)) model was trained and implemented with using scArches framework (Lotfollahi et al., 2022). Query organoid data was subsetted to RGCs and reference in vivo data (*10*) was subsetted to all RGC types (RG-vRG, RG-oRG, RG-tRG) from First_trimester and Second_trimester stages. Model was trained with 2 hidden layers and 30 latent dimensions using top 1500 highly variable genes using seurat_v3 flavor.

### Trajectory inference of organoid neuronal differentiation

Palantir (1.3.3) package was used perform trajectory analysis (*73*). Interneurons and Choroid plexus cells were excluded from the trajectory inference. Diffusion maps were calculated using top 5 Harmony corrected PCs and were used for multi-scale representation. A root cell from D50 dividing RGCs was selected, and a D90 IT neuron and a D90 non-IT neuron were chosen as terminal cells based on high expression of neuronal maturation genes MEF2C, CAMK2B and GRIN2B. Bifurcating trajectories were generated with *`run_palantir`* (500 waypoints, 30 nearest neighbours). Cells with high probability to belong to IT and DL/Non-IT branches were selected (q=.001, eps=.001). Gene expression and gene set score dynamics along neuronal differentiation and maturation were assessed by fitting a generalized additive model (GAM), using the *pyGAM* package (v0.9.1; Servén & Brummitt, 2018), to the IT and DL/Non-IT pseudotime trajectories, after excluding radial glial cells (RGCs).

### Progenitor sorting

For progenitor sorting of mouse primary cortical cells, cells were dissociated at 2 days in vitro using the NeuroCult Enzymatic Dissociation Kit in accordance with the manufacturer’s instructions. Dissociated cells were incubated with magnetic beads conjugated with anti-mouse prominin-1 (anti-mouse CD133, Miltenyi Biotec, Cat #130-092-564) in MACS buffer (Miltenyi Biotec, Cat #130-091-376 and #130-091-222) at 4°C for 15 min.

CD133-positive selection was carried out with LS columns (Miltenyi Biotec, Cat #130-042-401) according to the manufacturer’s instructions.

For progenitor sorting of human cortical organoid-derived cells, two days before, organoids were dissociated using Papain Dissociation System (Worthington Biochemical Corporation) and plated in Matrigel-coated 6-well plates in Nb/N2B27 medium. For MACS, cells were dissociated using the NeuroCult Enzymatic Dissociation Kit in accordance with the manufacturer’s instructions. Dissociated cells were incubated with APC-conjugated anti-CXCR4 (anti-CD184, Miltenyi Biotec, Cat #130-100-070) in MACS buffer at 4°C for 10 min. Then, cells were washed and incubated with anti-APC microbeads (Miltenyi Biotec, Cat #130-100-070) at 4°C for 15 min. CD184-positive selection was carried out with LS columns according to the manufacturer’s instructions.

For progenitor sorting of 2D human cortical cells, cells were dissociated using NeuroCult Enzymatic Dissociation Kit following manufacturer’s instructions. Dissociated cells were incubated with magnetic beads conjugated anti-human prominin-1 (anti-human CD133, Miltenyi Biotec, Cat #130-097-049) with FcR Blocking Reagent in MACS buffer at 4°C for 30 min. CD133-positive selection was carried out with LS columns according to the manufacturer’s instructions.

The sorted cells were resuspended in MiR05 buffer (Oroboros Instruments) and used for measuring high-resolution monitoring of respiratory activity using Oxygraph-2k (Oroboros Instruments).

### Oxygraph

Depending on the species and cell birthdate, MiR05 buffer was added to adjust to 6,000,000-80,000,000 cells/ml. One hundred microlitres of this cell suspension were injected per chamber of the oxygraphy O2K, with subsequent injections of the following compounds in a specific order until reaching the final concentration, depending on the purpose.

For measuring maximum OXPHOS (phosphorylating) and ETC (non-phosphorylating) capacity: (1) F-catalase (Merck, Cat#C9322), 556 U/ml; (2) digitonin (Merck, Cat#300410), 10 μg/ml; (3) pyruvate (Merch P2256), 5 mM; (4) malate (Merck, Cat#02288), 0.5 mM; (5) ADP (Calbiochem, Cat#117105), 1 mM; (6) glutamate (Merck, Cat#G1251), 10 mM; (7) succinate (Merck, Cat#S2378), 10 mM; (8) CCCP (Merck, Cat#C2759), Δ1 μM until maximum respiration reached; (9) rotenone (Merck, Cat#R8875), 75 nM; (10) glycerophosphate (Merck, Cat#G7886), 10 mM; (11) antimycin A (Merck, Cat#A8674), 250 nM; (12) sodium ascorbate (Merck, Cat#A7631), 2 mM; (13) N,N,N^′^,N^′^-Tetramethyl-p-phenylenediamine dihydrochloride (TMPD) (Merck, Cat#T3134) 0.5 mM; (14) cytochrome C (Merck, Cat#C7752), 10 μM; (15) sodium azide (Merck, Cat#S2002), 100 mM.

For measuring fatty acid-dependent OXPHOS capacity:

(1) F-catalase, 556 U/ml; (2) digitonin,10 μg/ml; (3) malate, 0.1 mM; (4) ADP, 1 mM; (5) octanoylcarnitine (TOCRIS, Cat #0605), 40 µM; (6) malate, 0.4 mM; (7) pyruvate, 5mM; (8) glutamate, 10mM; (9) succinate, 10mM; (10) CCCP, Δ1μM until maximum respiration reached; (11) rotenone, 75nM; (12) antimycin A, 250nM.

Oxygen saturation was maintained ≥ 40μM throughout the experiment, and between 170 and 200μM before the measurement of isolated complex IV (CIV) activity.

The reported values of maximum OXPHOS (phosphorylating), maximum ETC (non-phosphorylating), and fatty acid-dependent OXPHOS were calculated as succinate minus antimycin A (non-mitochondrial respiration), CCCP minus antimycin A, and octanoylcarnitine minus antimycin A, respectively.

### Lipid uptake assay

For measuring lipid uptake, the culture medium was replaced with fresh medium containing 5 µM BODIPY FL C12 (Invitrogen, Cat #D3822), conjugated to BSA in a 2:1 ratio. Cells were incubated with BODIPY for 15 min at 37°C in the dark. Cells were washed and dissociated using the NeuroCult Enzymatic Dissociation Kit. For antibody staining, 500,000 cells were resuspended in 100 µl of antibody-mixed FACS buffer (1% FBS and 1 mM EDTA in PBS). For mouse cells, anti-CD133 (Prominin-1) monoclonal antibody (1:50, Invitrogen, Cat #17-1331-81). For human cells, anti-CD184 (CXCR4) monoclonal antibody (1:100, Invitrogen, Cat #17-9991-82) and anti-CD171 monoclonal antibody (1:25, Invitrogen, Cat #12-1719-42). After 20 min incubation at 4°C, cells were washed twice and resuspended in 200 µl of FACS buffer. BODIPY fluorescence intensity and antibody fluorescence were measured using a 4-laser BD FACSymphony™ A1 flow cytometer with BD FACSDiva Software (version 9.0.2), and subsequently analyzed using BD FlowJo v10.10.0 (Becton Dickinson & Company).

### Acetyl-CoA measurement using liquid-chromatography mass spectrometry (LC-MS)

To measure acetyl-CoA, mouse primary cultures and human cortical organoid cultures were snapfrozen after 48h and lysed using an ice-cold 10% trichloroacetic acid solution. Subsequently, the samples were purified using and Oasis HLB SPE column (Waters). Dried acetyl-CoA samples were then resuspended in 50µl of 5% sulfosalicylic acid before loading on the LC-MS. A Dionex UltiMate 3000 LC System with a thermal autosampler set at 4 °C, coupled to a Q-Exactive Orbitrap MS (Thermo Scientific) was used and 10 μL of each sample was injected on a C18 column (Acquity UPLC HSS T3, 1.8 µm, 2.1 × 100 mm). The separation of metabolites was achieved at 40 °C with a flow rate of 0.25 mL/min. A gradient was applied for 40 minutes (A, 10 mM tributyl-amine and 15 mM acetic acid; B, methanol) to separate the targeted metabolites (0 min at 0% B, 2 min at 0% B, 7 min at 37% B, 14 min at 41% B, 26 min at 100% B, 30 min at 100% B, 31 min at 0% B; and 40 min at 0% B. The MS operated in negative full-scan mode (m/z range 70–500 and 190–300 from 5–25 min) using a spray voltage of 4.9 kV, capillary temperature of 320 °C, sheath gas at 50.0 and auxiliary gas at 10.0. Data were analysed using the Xcalibur software (Thermo Scientific) and normalized by internal standard d27 myristic acid and protein content.

### Statistical analysis

All statistical analyses were performed using GraphPad Prism 10 (GraphPad Software) or SigmaPlot 12.5 (Systat Software Inc). A Kolmogorov-Smirnov test or Shapiro-Wilk normality test was used to test the normality. Parametric data were analyzed by t-test, one-way or two-way ANOVA followed by Dunnett’s, or Tukey’s post hoc analysis for comparisons of multiple samples. Non-parametric data were analyzed by the Mann-Whitney test followed by Dunn’s post hoc analysis for comparisons of multiple samples. P values < 0.05 were considered statistically significant. Data are presented as mean ± SEM or median.

---

**Fig. S1.**
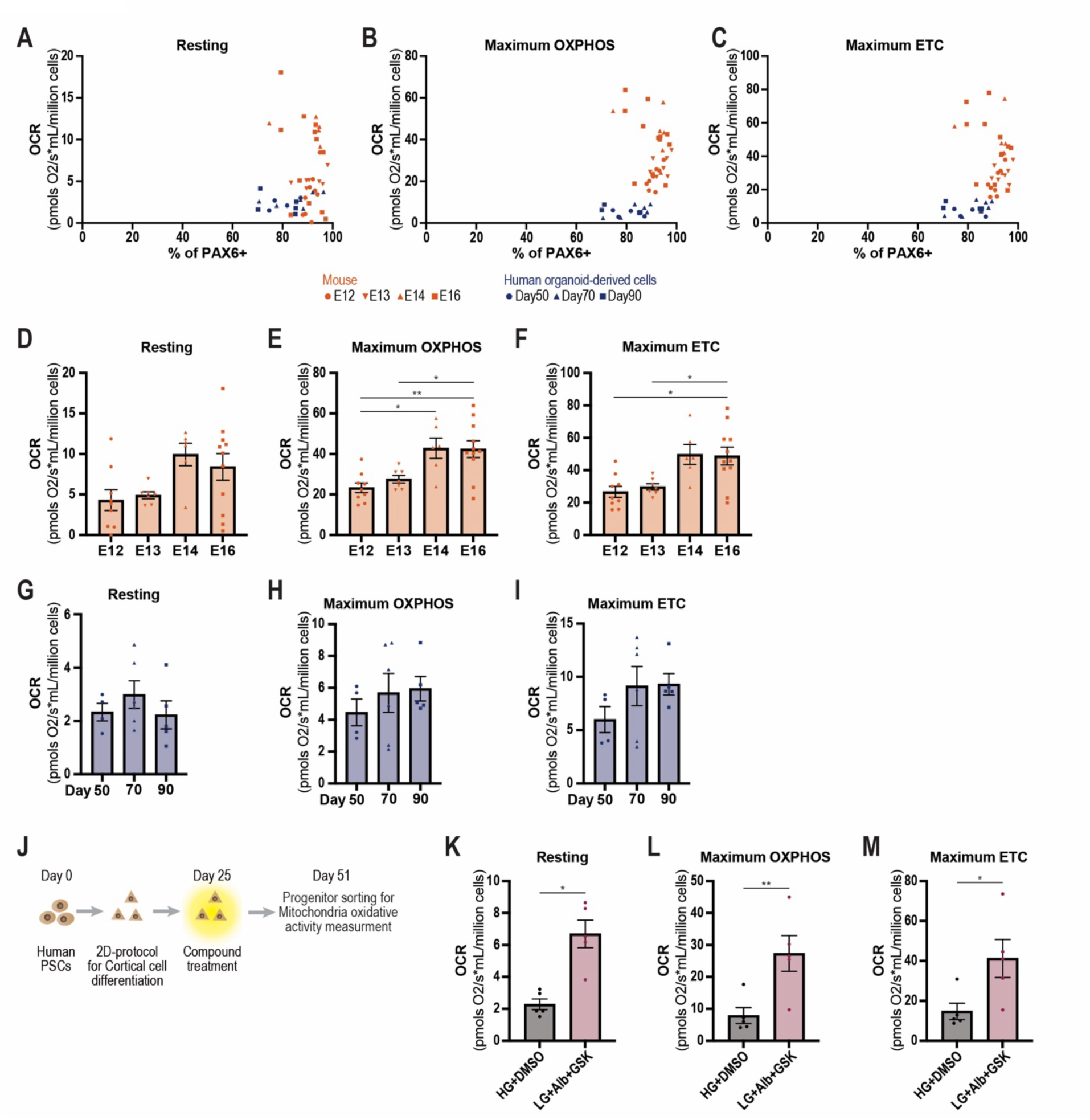
Comparison of oxygen consumption rate in cortical cells over time, across species, and under different conditions. (A,B,C) Quantified proportion of progenitor marker PAX6-positive and relation with (G) resting OCR, (H) maximum OXPHOS, and (I) maximum ETC. Each data point represents an individual experiment. (D-I) Quantification of oxygen consumption rate (OCR) cortical progenitors at different developmental stages. Each data point represents an individual experiment. (D-F) Mouse primary progenitors (derived from embryonic days (E)12, 13, 14, 16. N=9, 7, 6, 11 experiments). (G-I) Human cortical organoid-derived progenitors (derived from differentiation day 50, 70, 90. N=4,6, 5 different batches respectively). Dunnett’s or Dunn’s multiple comparisons test. (A, D) Resting. (B, E) Maximum oxidative phosphorylation (OXPHOS) capacity under coupled conditions. (C, F) Maximum electron transport chain (ETC) capacity under uncoupled condition (J) Schematic of timeline of mitochondria oxidative activity measurement using 2D-differentiation protocol-derived human cortical progenitors. (K, L, M) Quantification of OCR following different chemical compound treatments on cortical progenitors at day 51. Each data point represents an individual experiment from five independent differentiation batches. High glucose (25 mM) with DMSO (HG+DMSO: N=5 different batches). Low glucose (2.5 mM) with 0.5% AlbuMAX and 5µM GSK 2837808A (LG+Alb+GSK: N=5 different batches). Paired t test. (K) Resting. (L) Maximum OXPHOS. (M) Maximum ETC. *P < 0.05, **P < 0.01

**Fig. S2.**
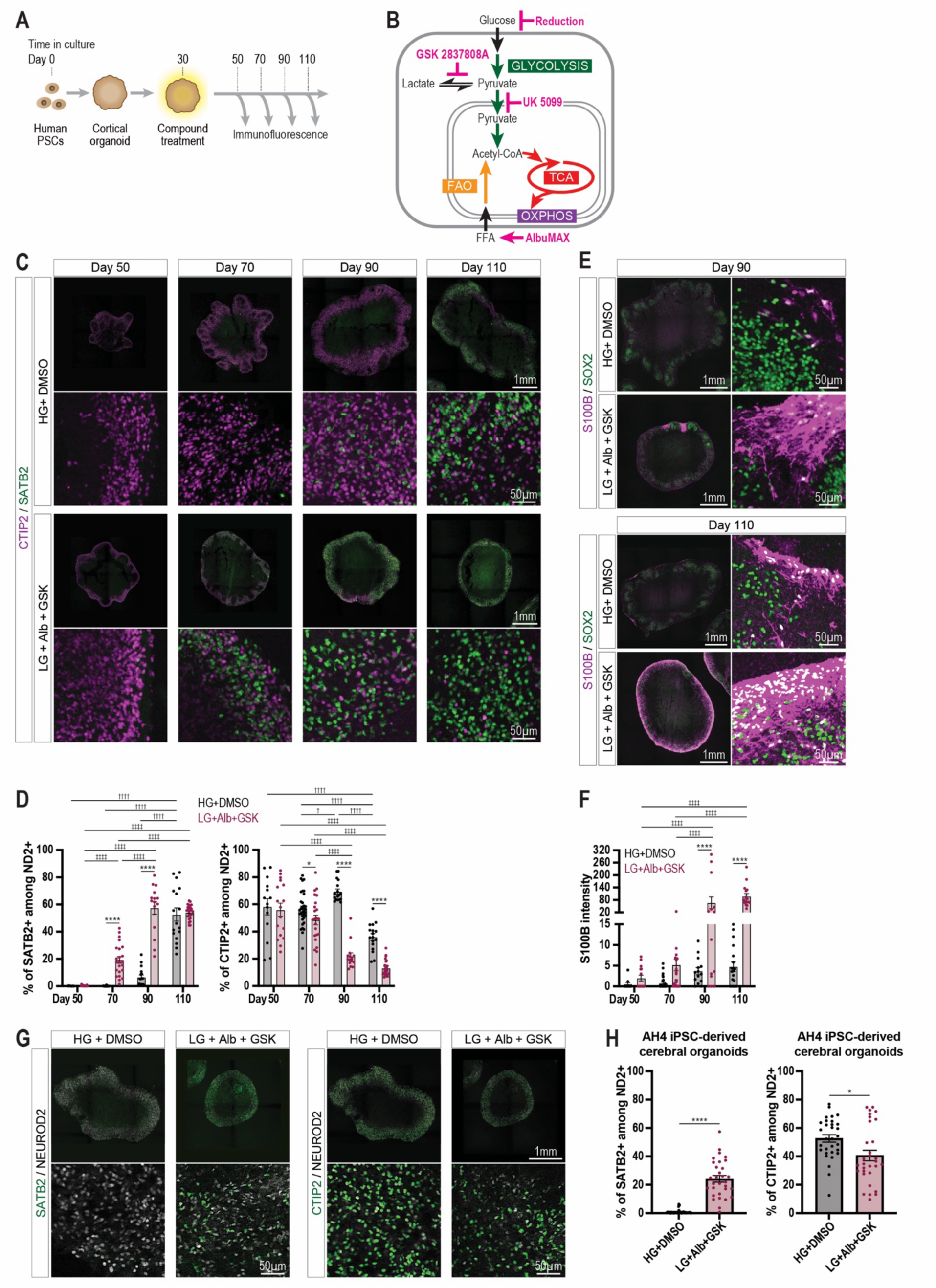
Increased mitochondrial metabolic rates accelerate human corticogenesis. (A) Schematic of timeline for human cortical organoid experiments. (B) Metabolic pathways targeted by indicated chemical compounds. Tricarboxylic acid cycle (TCA). Oxidative phosphorylation (OXPHOS). (C) Representative images of cortical organoid for SATB2 and CTIP2. High glucose (25 mM) with DMSO (HG+DMSO) (top) and Low glucose (2.5 mM) with 0.5% AlbuMAX and 5µM GSK 2837808A (LG+Alb+GSK) (bottom). (D) Quantification of the proportion of (Left) SATB2 or (Right) CTIP2-positive cells. Each data point represents an individual organoid. High glucose (25 mM) with DMSO (HG+DMSO): differentiation day 50, 70, 90, 110. N=14, 34, 17, 17 organoids obtained from five independent differentiation batches. Low glucose (2.5 mM) with 0.5% AlbuMAX and 5µM GSK 2837808A (LG+Alb+GSK): differentiation day 50, 70, 90, 110. N=17, 23, 15, 24 organoids organoids were obtained from five independent differentiation batches. Tukey’s multiple comparisons test. (E) Representative images of cortical organoid. S100B is used for labelling astroglial cells. (F) Quantification of mean signal intensity of S100B from each organoid. Each data point represents an individual organoid. High glucose with DMSO (HG+DMSO): differentiation day 50, 70, 90, 110. N=16, 30, 14, 18 organoids from five independent differentiation batches. Low glucose with 0.5% AlbuMAX and 5µM GSK 2837808A (LG+Alb+GSK): differentiation day 50, 70, 90, 110. N=15, 23, 14, 16 organoids). Tukey’s multiple comparisons test. (G) Representative images of cortical organoid derived from a human iPSC line, AH4. (Left) SATB2 (Right) CTIP2. (H) Quantification of the proportion of (Left) SATB2 or (Right) CTIP2-positive cells. Each data point represents an individual organoid. High glucose (25 mM) with DMSO (HG+DMSO): N=31 organoids from three independent differentiation batches. Low glucose (2.5 mM) with 0.5% AlbuMAX and 5µM GSK 2837808A (LG+Alb+GSK): N=31 organoids from three independent differentiation batches. Mann-Whitney test. * † ‡P < 0.05, ** †† ‡‡P < 0.01, *** ††† ‡‡‡P < 0.001, **** †††† ‡‡‡‡P < 0.0001.

**Fig. S3.**
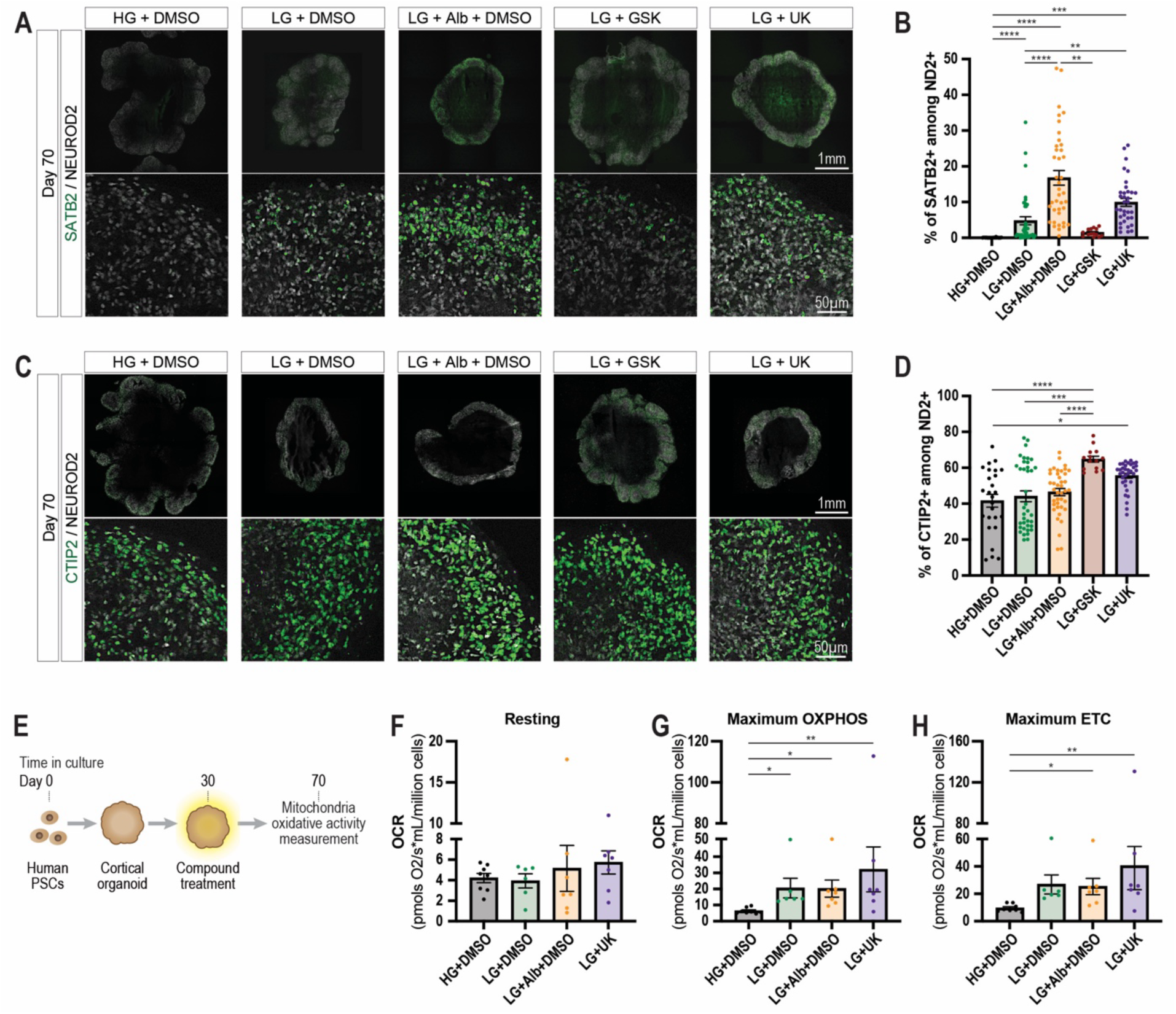
FAO is the main metabolic pathway accelerating temporal patterning of human corticogenesis. (A, C) Representative images of cortical organoid at day 70. (A) SATB2. (C) CTIP2. (B) Quantification of the proportion of SATB2-positive cells. Each data point represents an individual organoid. High glucose with DMSO HG+DMSO: N=26 organoids from four independent differentiation batches. Low glucose with DMSO (LG+DMSO): N=40 organoids from four independent differentiation batches. Low glucose with 0.5% AlbuMAX and DMSO LG+Alb+DMSO: N=39 organoids from four independent differentiation batches. Low glucose with GSK 2837808 (LG+GSK: N=14 organoids from four independent differentiation batches. Low glucose with 1.25µM UK 5099 (LG+UK): N=34 organoids from four independent differentiation batches. Dunn’s multiple comparisons test. (D) Quantification of the proportion of CTIP2-positive cells. Each data point represents an individual organoid. High glucose with DMSO. HG+DMSO: N=26 organoids). Low glucose with DMSO (LG+DMSO: N=38 organoids from four independent differentiation batches. Low glucose with 0.5% AlbuMAX and DMSO (LG+Alb+DMSO): N=40 organoids from four independent differentiation batches. Low glucose with GSK 2837808 (LG+GSK): N=14 organoids from four independent differentiation batches. Low glucose with 1.25µM UK 5099 (LG+UK: N=34 organoids) Dunn’s multiple comparisons test. (E) Schematic of timeline of mitochondria oxidative activity measurement using human cortical organoids under different chemical compound treatment conditions. (F, G, H) Quantification of OCR at different chemical compound treatments on human cortical organoid-derived cortical cells at day 70. Each data point represents an individual experiment from eight independent differentiation batches. High glucose with DMSO (HG+DMSO: N=8 experiments). Low glucose with DMSO (LG+DMSO: N=6 experiments). Low glucose with 0.5% AlbuMAX and DMSO (LG+Alb+DMSO: N=7 experiments). Low glucose with 1.25µM UK 5099 (LG+UK: N=7 experiments) Dunn’s multiple comparisons test. (F) Resting. (G) Maximum OXPHOS. (H) Maximum ETC. *P < 0.05, **P < 0.01, ***P < 0.001, ****P < 0.0001.

**Fig. S4.**
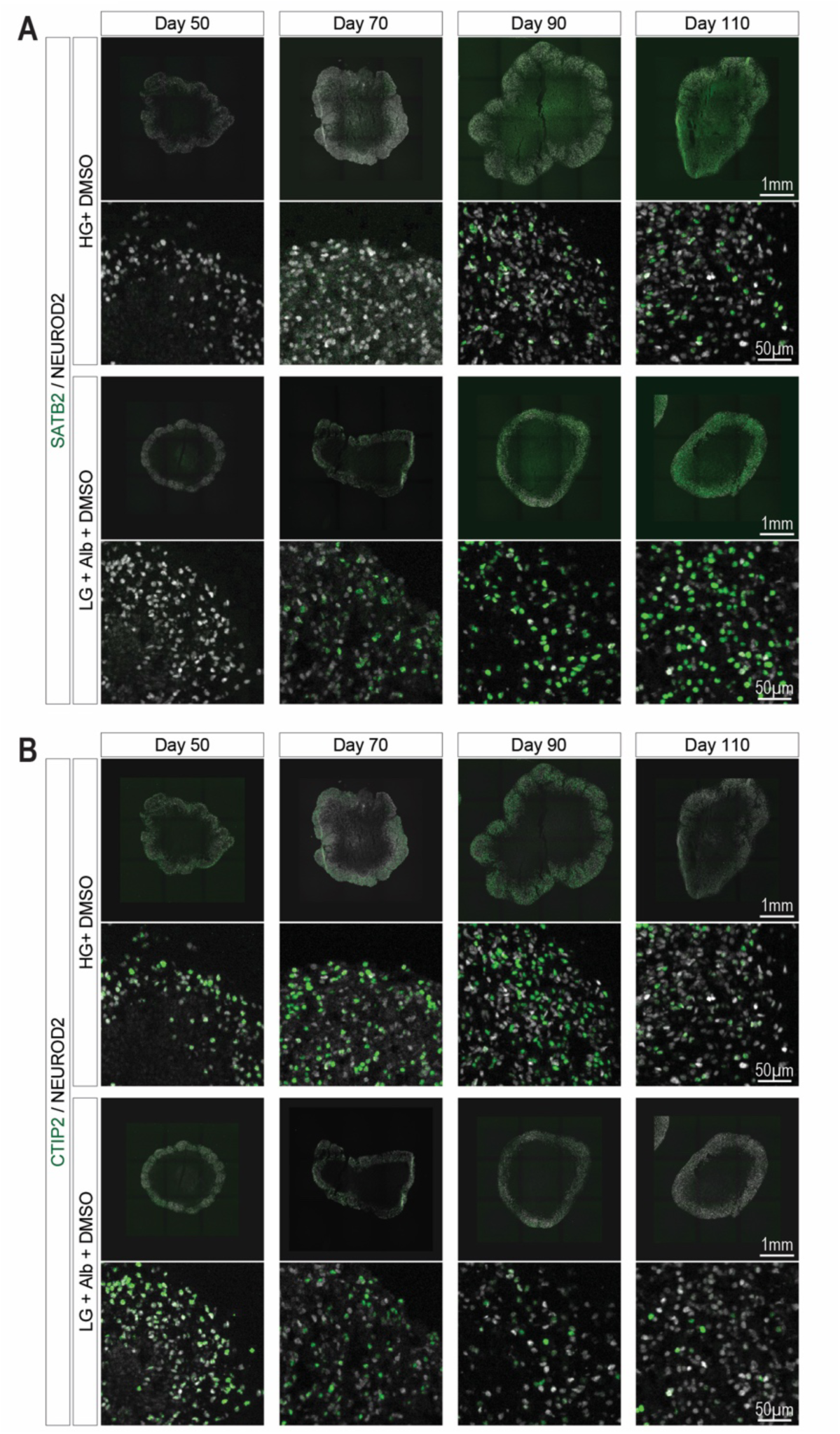
Developmental dynamics of layer marker expression in AlbuMAX-treated organoids. (A, B) Representative images of cortical organoid at day 70. (A) SATB2. (B) CTIP2. Organoid images of SATB2 and CTIP2 are derived from the same section.

**Fig. S5.**
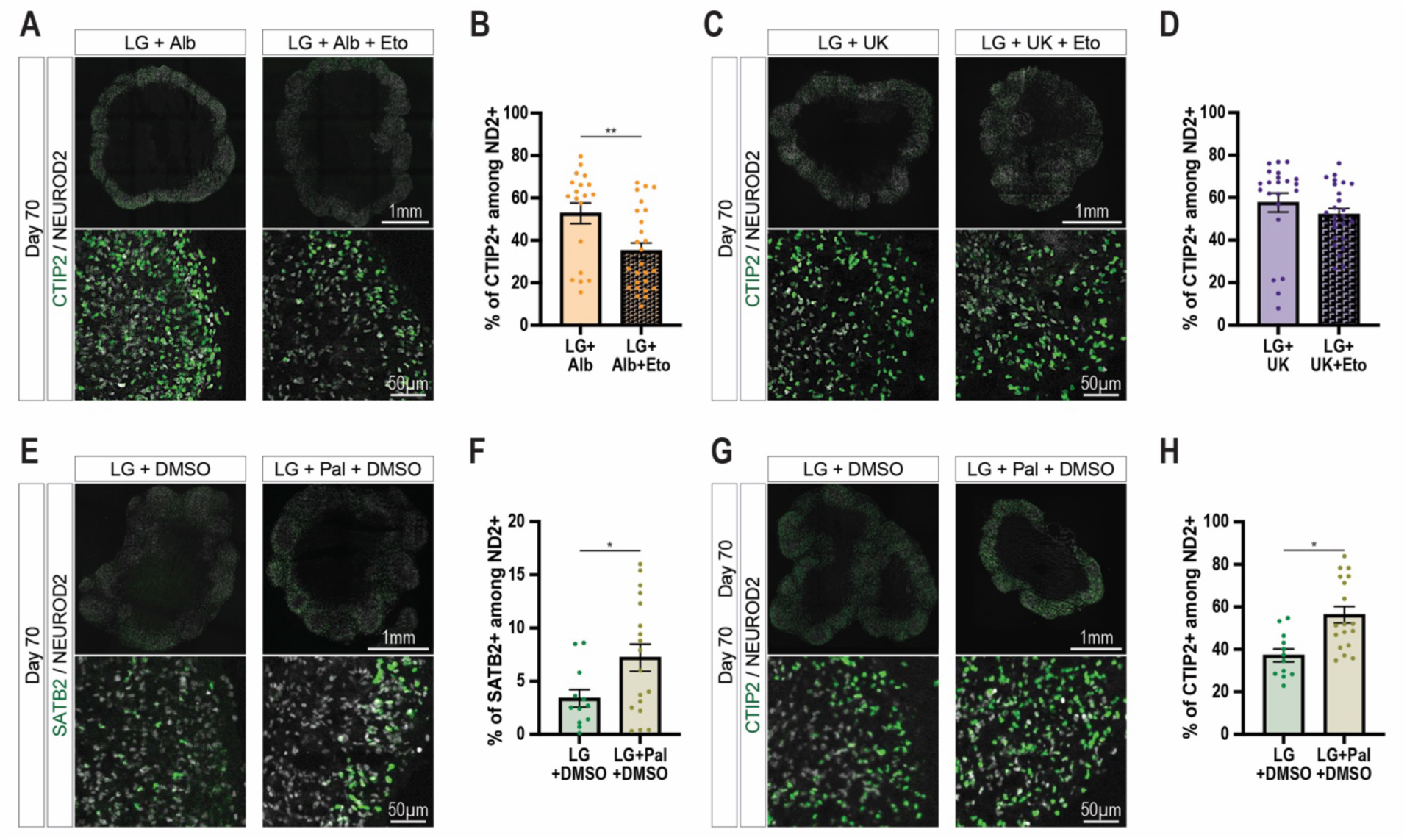
Impact of fatty acid oxidation metabolism on layer marker expression. (A, C, E, G) Representative images of cortical organoid at day 70. (A, C, G) CTIP2. (E) SATB2. (B) Quantification of the proportion of CTIP2-positive cells. Each data point represents an individual organoid. Low glucose with 0.5% AlbuMAX and DMSO (LG+Alb): N=19 organoids from three independent differentiation batches. Low glucose with 0.5% AlbuMAX and 50µM Etomoxir (LG+Alb+Eto): N=26 organoids from three independent differentiation batches. Mann-Whitney test. (D) Quantification of the proportion of CTIP2-positive cells. Each data point represents an individual organoid. Low glucose with 1.25µM UK 5099 (LG+UK): N=22 organoids from three independent differentiation batches. Low glucose with 1.25µM UK 5099 and 50µM Etomoxir (LG+UK+Eto): N=25 organoids from three independent differentiation batches. Mann-Whitney test. (F, H) Quantification of the proportion of (F) SATB2 or (H) CTIP2-positive cells. Each data point represents an individual organoid. Low glucose with DMSO (LG+DMSO): N=12 organoids from two independent differentiation batches. Low glucose with 100µM Palmitate and DMSO (LG+Pal+DMSO): N=18 organoids from two independent differentiation batches. (F) Unpaired t test. (H) Mann-Whitney test. *P < 0.05, **P < 0.01

**Fig. S6.**
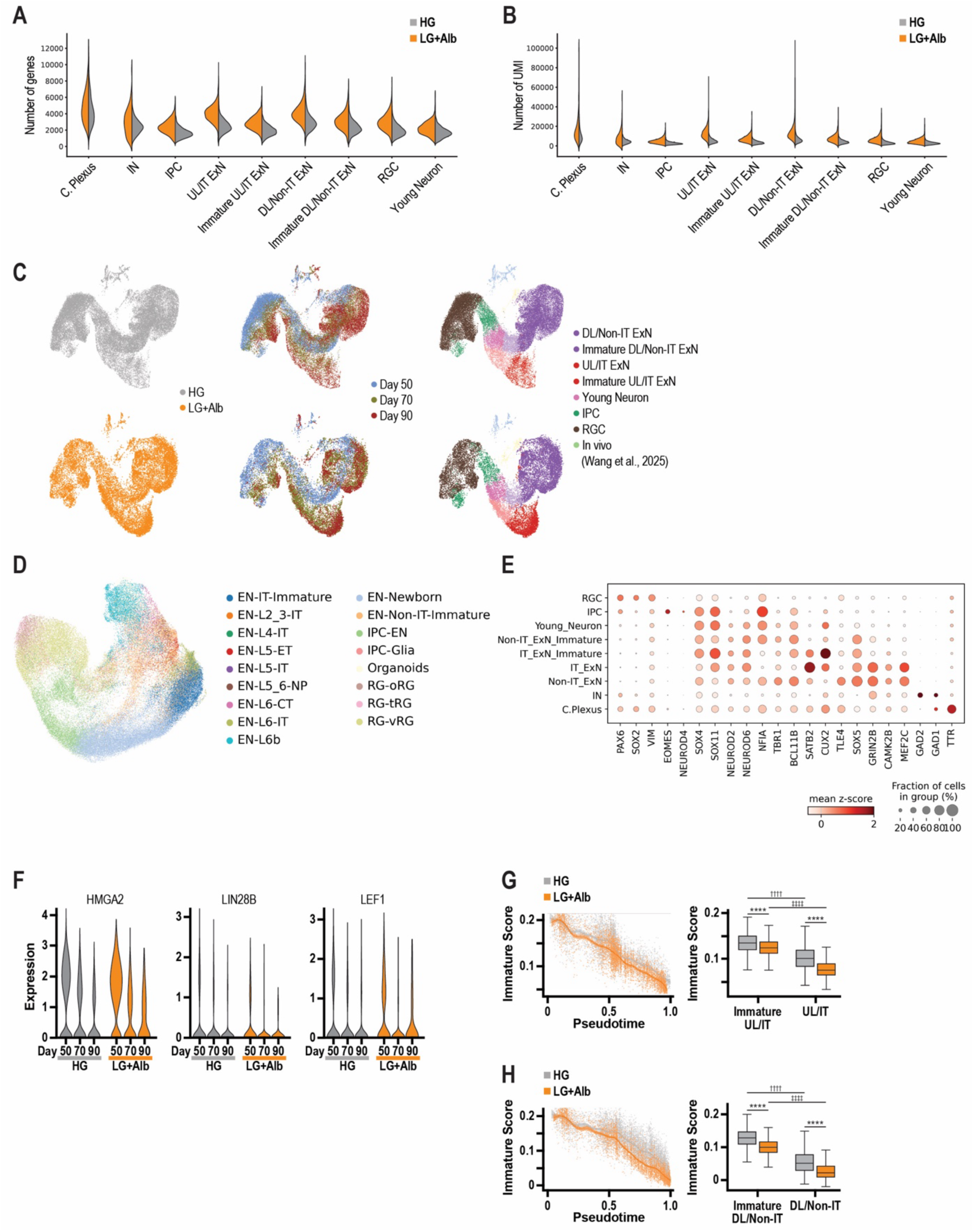
SnRNAseq analysis of cortical organoids following Alb treatments. (A) Violin plot of number of genes detected per cell type. (B) Violin plot of number of total UMI (unique molecular identifier) per cell type. (C) UMAP of High glucose (25 mM) with DMSO (HG) (Top) and Low glucose with 0.5% AlbuMAX and DMSO (LG+Alb) treated organoids (Bottom) after Harmony integration. (D) UMAP plot of in vivo fetal cortex and organoid cells showing cell type annotations for the in vivo dataset after Harmony integration. (E) Dot plot showing the expression of known cell type-specific marker genes. Dot colour intensity represents the mean z-score of the gene, and dot size indicates the percentage of cells expressing the gene within the cell type. (F) Violin plot showing normalized expression of differentially expressed canonical early RGC markers. (F) (Left) Generalized additive model (GAM) fit of immature IT neuron score along control and FFA-treated IT neuron trajectories. (Right) Boxplot of immature IT score over immature IT and IT neurons across control and FFA treatment. Mann-Whitney test two-sided with Bonferroni correction. (H) (Left) Generalized additive model (GAM) fit of immature non-IT neuron score along control and FFA-treated IT neuron trajectories. (Right) Boxplot of immature non-IT score over immature non-IT and non-IT neurons across control and FFA treatment. * † ‡P < 0.05, ** †† ‡‡P < 0.01, *** ††† ‡‡‡P < 0.001, **** †††† ‡‡‡‡P < 0.0001.

**Fig. S7.**
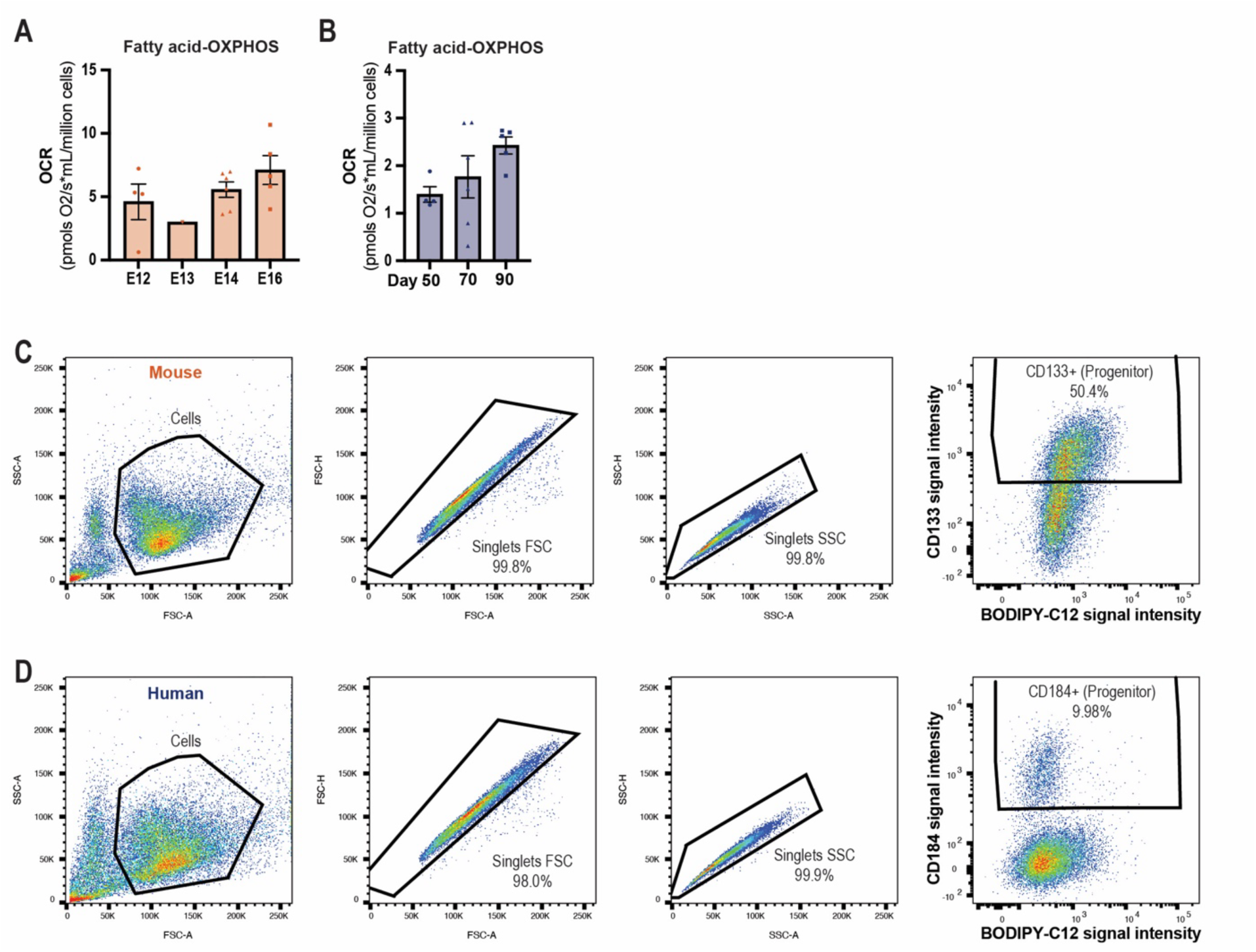
Comparison of FAO activity and fatty acid uptake between mouse and human cortical progenitors. (A) Quantification of fatty acid treatment-induced OCR across various developmental stages. Each data point represents an individual experiment. Mouse primary progenitors (derived from embryonic days (E)12, 13, 14, 16. N=4, 1, 6, 5 experiments). Kruskal-Wallis test. (B) Quantification of fatty acid treatment-induced OCR across various developmental stages. Each data point represents an individual experiment. Human organoid-derived progenitors (derived from differentiation day 50, 70, 90. N=4, 6, 5 different batches). Kruskal-Wallis test. (C, D) Representative flow cytometry plots and gating strategy for measuring BODIPY intensity in (C) mouse and (D) human cortical cells.

**Fig. S8.**
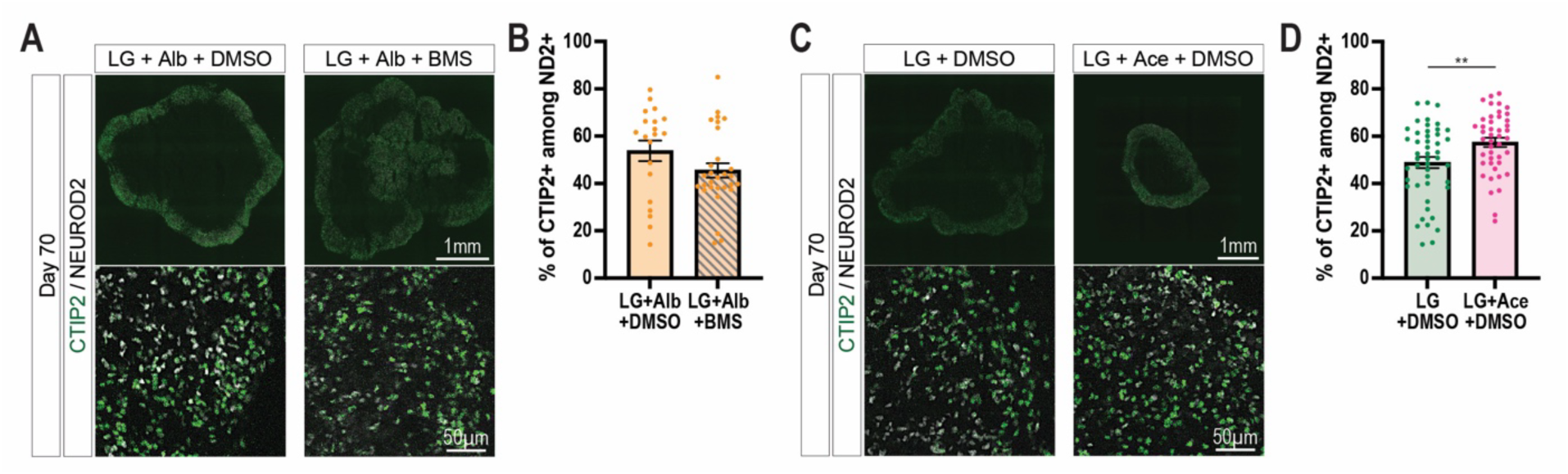
Impact of acetate and ACLY inhibitor treatments on layer marker expression. (A, C) Representative images of cortical organoid at day 70. CTIP2. (B) Quantification of the proportion of CTIP2-positive cells. Each data point represents an individual organoid. Low glucose with 0.5% AlbuMAX and DMSO (LG+Alb: N=20 organoids from three independent differentiation batches. Low glucose with 0.5% AlbuMAX and 5µM BMS (LG+Alb+BMS 303141: N=29 organoids from three independent differentiation batches. Mann-Whitney test. (D) Quantification of the proportion of SATB2-positive cells. Each data point represents an individual organoid. Low glucose with DMSO LG+DMSO: N=47 organoids from five independent differentiation batches. Low glucose with 100 µM Acetate and DMSO . LG+Ace+DMSO: N=44 organoids from five independent differentiation batches. Mann-Whitney test. **P < 0.01.

